# Persistence of learning-induced synapses depends on neurotrophic-priming of glucocorticoid receptors

**DOI:** 10.1101/623389

**Authors:** M. Arango-Lievano, A. Borie, Y. Dromard, M. Murat, M.G. Desarménien, M.J. Garabedian, F. Jeanneteau

## Abstract

Stress can either promote or impair learning and memory. Such opposing effects depend on whether synapses persist or decay after learning. Maintenance of new synapses formed at the time of learning upon neuronal network activation depends on the stress hormone activated glucocorticoid receptor (GR) and neurotrophic factor release. Whether and how concurrent GR and neurotrophin signaling integrate to modulate synaptic plasticity and learning is unknown. Here we show that deletion of the neurotrophin BDNF-dependent GR phosphorylation sites (GR-PO_4_) impairs long-term memory retention and maintenance of newly formed postsynaptic dendritic spines in the mouse cortex after motor skills training. Chronic stress and the BDNF polymorphism Val66Met disrupt the BDNF-dependent GR-PO_4_ pathway necessary for preserving training-induced spines and previously acquired memories. Conversely, enrichment living promotes spine formation but fails to salvage training-related spines in mice lacking BDNF-dependent GR-PO_4_ sites, suggesting it is essential for spine consolidation and memory retention. Mechanistically, spine maturation and persistence in the motor cortex depend on synaptic mobilization of the glutamate receptor GluA1 mediated by GR-PO_4_. Together, these findings indicate that regulation of GR-PO_4_ via activity-dependent BDNF signaling is important for learning-dependent synapses formation and maintenance. They also define a new signaling mechanism underlying these effects.

**SIGNIFICANCE STATEMENT:** Signal transduction of receptors tyrosine kinase and nuclear receptors is essential for homeostasis. Phosphorylation is one of the currencies used by these receptors to support homeostatic reactions in learning and memory. Here we show that consolidation of learning-induced neuroplasticity is made possible via stress activated glucocorticoid nuclear receptor phosphorylation through the brain-derived neurotrophic tyrosine kinase pathway. Crosstalk between these pathways is specific of cell types and behavioral experience (*e.g*. learning, stress and enrichment living). Disruption of this response may contribute to the pathophysiology of stress-related disorders and treatment resistance.

## INTRODUCTION

Stress modifies adaptive behaviors such as learning and memory (1). Glucocorticoids are a stress hormone that signal via the glucocorticoid receptor (GR) and can either promote or impair learning and memory (2). Whereas prolonged secretion of glucocorticoids during chronic stress disrupts learning and memory, its release at the time of learning may promote it (3). An acute rise in glucocorticoid levels at the time of learning stimulates the formation and stabilization of new dendritic spines, and eliminates synapses established before learning. New dendritic spines require protein synthesis, which is initiated in the neuronal networks targeted by behavioral experience (4–6). For example, motor learning instructs remodeling of dendritic spines in excitatory neurons of the motor cortex (7–9). Stabilization of new spine synapses forges new learning-induced connectivity that correlates with memory consolidation (10, 11). However, the pathways and molecular mechanisms affecting spine stabilization during learning and memory *in vivo* are not understood.

Like glucocorticoids via GR, brain derived neurotrophic factor (BDNF) through its receptor TrkB stabilizes newly formed synapses and fosters learning and memory (12–14). BDNF is also critical for modulating the impact of stress in the corticolimbic and mesolimbic systems (15, 16). Behavioral actions of glucocorticoids and BDNF are complementary, and play roles in avoidance, fear, coping, and impulse control (17, 18). The influence on GR by the BDNF pathway likely relies on activity-dependent release of BDNF, which is reduced in a BDNF genetic variant Val66Met associated with impaired response to stress (19–21). BDNF signaling through TrkB alters the GR transcriptome through changes in GR phosphorylation (GR-PO_4_) and can affect neuronal plasticity (22–24). However, it remains unclear whether BDNF-dependent GR-PO_4_ mediates the persistence of new spines associated with learning and memory, and if activity-dependent secretion of BDNF is also required.

Here we used two-photon in vivo microscopy of learning-associated dendritic spine remodeling to examine the effects of BDNF-dependent GR-PO_4_ pathway in a newly developed GR-PO_4_ deficient mouse and in a mouse carrying the Val66Met polymorphism of BDNF. We found that GR-PO_4_ and BDNF secretion were both important for the formation and maintenance of new spines after learning through the synaptic recruitment of glutamate receptor GluA1. Our findings uncover an important mechanism for how acute glucocorticoids can direct cell-type specific response to store and retain new information upon learning. By turning off this mechanism, chronic stress impaired cell-type specific contextual GR response.

## RESULTS

### Timing and specificity of BDNF-dependent GR-PO_4_ expression in motor cortex

The corticosterone-GR pathway is required for the acquisition of new motor skills (25), but its dependence on BDNF-dependent GR-PO_4_ is unknown. To unravel this, we first assessed whether learning might affect GR-PO_4_ using a rotarod learning paradigm. Mice were left untrained or trained for 2 days. Forty-five minutes after the training, both control untrained or trained mice were euthanized and expression in the cortex of GR-PO_4_ at the BDNF-dependent sites (S152 and S284 in mice correspond to S155 and S287 in rat numbering scheme as previously described (23, 26)), as well as c-FOS as an index of neuronal response were determined by immunohistochemistry. Training increased the level of GR-PO_4_ at S152 and S284 in motor areas of cortex compared to the untrained controls (Figure S1A). The thy1-YFP marked excitatory and parvalbumin (PV) inhibitory neurons are among the cell types exhibiting high levels of GR-PO_4_ in M1 primary motor cortex (Figure S1B). Both cells types displayed increased c-FOS protein abundance upon training. Training also raised the percentage of cells harboring BDNF-dependent GR-PO_4_ in all layers of the M1 cortex (Figure S1C). In fact, training also raised the proportion of cells double-labeled with GR-PO_4_ and c-FOS (Figure S1D), and increased GR-PO_4_ in PV and thy1 neurons (Figure S1E,F). Induction of GR-PO_4_ was site-specific (S152/S284 versus S234, Figure S1A) and did not persist 24 hours post-training, similar to c-FOS in cortical areas relevant for motor skill learning (Figure S1G). Therefore, GR-PO_4_ at BDNF-dependent sites increased with training in the motor cortex and is linked to neuronal c-FOS response.

### Deletion of GR-PO_4_ at BDNF-responding sites impaired motor skill retention

We next determined whether GR-PO_4_ at the BDNF-dependent sites affected learning and memory as well as plasticity of dendritic spines. To do this, we generated a conditional allele of GR consisting of exon 2 harboring serine to alanine mutations in the BDNF-dependent GR-PO_4_ sites S152A and S284A [sites S134 and S267 in human GR, and S155 and S287 in rat GR number (Figure 1A)]. Recombination into germ cells using the constitutive ROSA26-CRE mouse line resulted in the mutations stably transmitted to next generations (Figure S3). Cortical tissue from the GR phosphorylation deficient knockin mouse (which we term KI) showed no staining with GR-PO_4_ specific antibodies compared to littermate WT controls (Figure 1B). The same protein abundance of total GR was observed between the WT and KI mice (Figure S4). Likewise, mineralocorticoid receptor (MR) levels were unaffected in KI (Figure S4). Therefore, the KI mutant mice eliminated GR-PO_4_ without affecting the abundance of glucocorticoid receptors.

**Figure 1.**
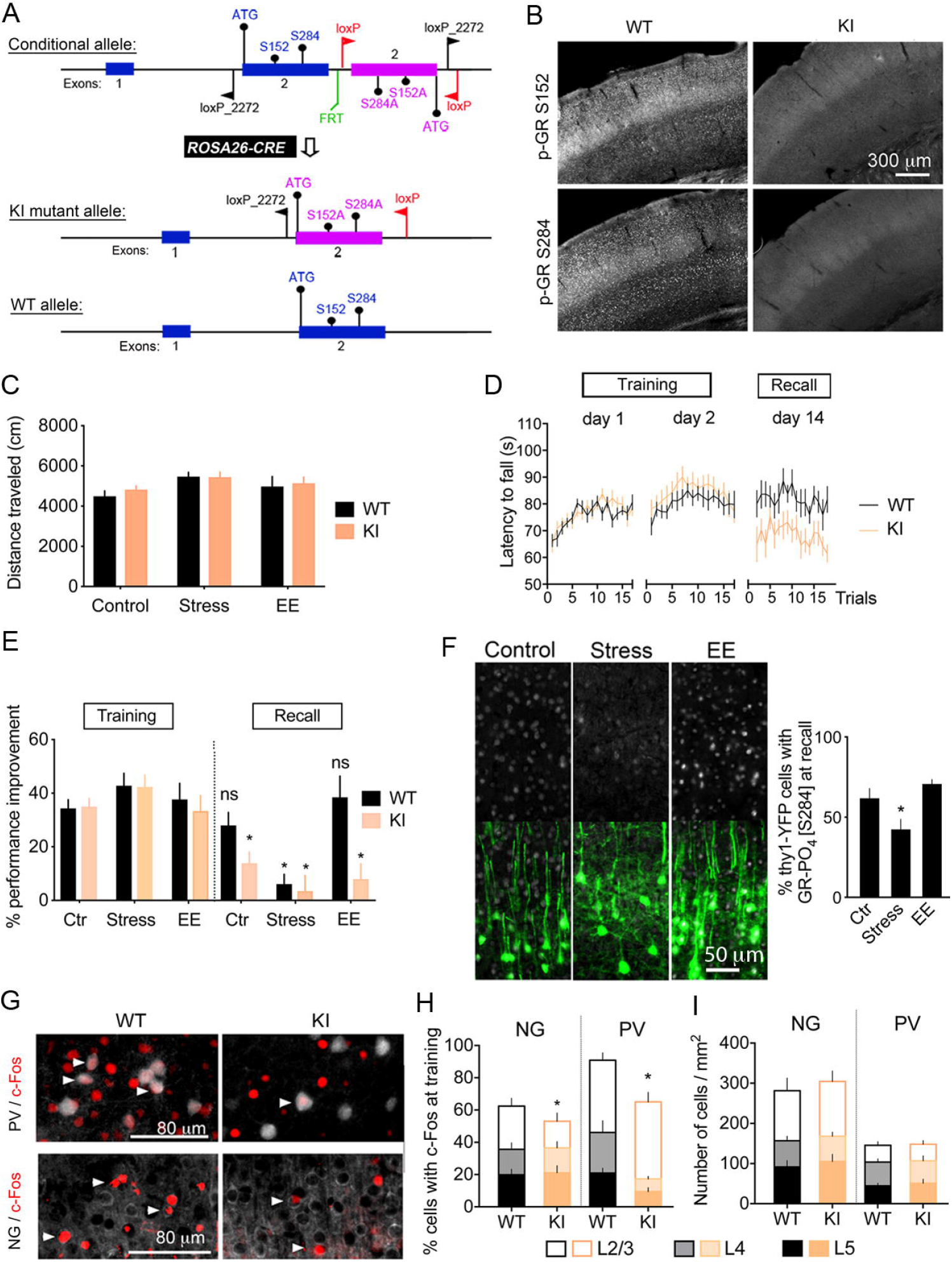
Deletion of BDNF-dependent GR-PO_4_ impairs retention of motor skill training. (A) Substitution of Ser152 and Ser284 (WT allele) by Ala152 and Ala284 (KI allele) in GR locus obtained by Cre-mediated recombination of loxP sites driven by the Rosa26 promoter. KI mice lack BDNF-dependent GR-PO_4_ sites. Details of the genetic constructs can be found in legend of figure S2 and methods. (B) Deletion of GR-PO_4_ immunostaining in KI homozygous mice compared to WT controls. (C) Spontaneous locomotion in an open field of mice reared in control conditions, chronic stress or enriched environment (EE) for 2 weeks starting immediately after the training. Means ±SEM of n=8 mice/ group, 2-way ANOVA: general effect of living conditions *F*_2,42_=3.43, *p*=0.039. (D) Motor skill learning monitored after 2 weeks of consolidation displayed as the latency to fall from the rotarod. Means ±SEM of n=8 mice/ group, 3-way ANOVA: Effect of genotype *F*_2,2_=3.075, *p*<0.05, post-hoc Tukey test: WT day 2 *p*=0.035, KI day 2 *p*=0.017; Effect on retention *F*_1,2_=11.33, *p*<0.05 post-hoc Tukey test: KI ^#^*p*=0.015. (E) Rotarod performance improvement to day 1 of mice reared in control, stress or EE conditions for 2 weeks after the training. Means ±SEM of n=8 mice/ group, 3-way ANOVA: Effect of retention *F*_1,84_=52.17, *p*<0.0001, genotype *F*_1,84_=9.1, *p*<0.01, stress x retention *F*_2,84_=7.8, *p*<0.001 and genotype x retention *F*_2,84_=6.55, *p*<0.05, post-hoc Tukey test between training and recall \**p*<0.05. (F) Effect of stress and EE between day 3 and 14 on GR-PO_4_ levels (grey) in thy1-YFP neurons (green) of M1 cortex. Means ±SEM of n=8 mice/ group. Unpaired t-test: *t*_(14)_=2.21 \**p*<0.05. See Figure S2 for larger fields of view. (G) Expression of *c-FOS* 45 min after 2 days of training in L2/3 neurogranin principal cells (NG) and L5 parvalbumin (PV) neurons of M1. Arrowheads point to co-expression. See Figure S2 for magnified pictures. (H) Means ±SEM of n=5 mice/ group, effect of genotype: 3-way ANOVA *F*_1,48_=8.33, *p*=0.0058, effect of cell type *F*_1,48_=10.95, *p*=0.0018, effect in cortical layers *F*_2,48_=31.24, *p*<0.0001 post-hoc Tukey test for NG cells L2/3 \**p*=0.026, L4 *p*=0.93 and L5 *p*=0.77; for PV cells L2/3 *p*=0.58, L4 \**p*=0.004 and L5 \**p*=0.046. (I) Number of NG and PV neurons/mm^2^ in M1. Means ±SEM of n= 5 mice/ group.

Deletion of the BDNF-dependent GR-PO_4_ sites resulted in adult mice of normal appearance and body weight under standard, stressed or enriched living conditions. There was no significant difference between the KI and WT mice in their degree of anxiety (as measured by thigmotaxis and elevated maze) (Figure S5A,B), despair behaviors (as assessed by tail suspension and forced swim tests)(Figure S5C,D) and locomotor activity when reared in standard living conditions as well as in stressful or in enriched environmental conditions (Figure 1C), previously showed to activate the corticosterone-GR pathway (27). KI mice exhibited normal learning abilities of new motor skills on the rotarod but impaired retention of the task compared to WT littermates (Figure 1D). Retention of new rotarod motor skills was also impaired in WT mice if exposed to chronic unpredictable stress immediately after training during the consolidation period, whereas enrichment living had no impact (Figure 1E). This is consistent with the reduction of BDNF-dependent GR-PO_4_ in motor cortex of mice reared in chronic stress (Figure 1F). What’s more, training-induced c-FOS expression in the excitatory and inhibitory (PV) neurons was reduced in KI mice compared to WT controls (Figure 1H,I). The effect of the KI showed in specific layers of the cortex differently whether PV or NG cells, suggesting putative inter-layers network compensation in KI mice. Therefore, GR-PO_4_ disruption impaired activation of M1 cortex as determined by c-FOS expression, and motor skill retention, which depend on BDNF and corticosterone (25, 28).

### GR-PO_4_ is required for the maintenance of new dendritic spines formed at training

Previous studies have shown that *de novo* spine formation and maintenance contribute to the storage of new motor skills by creating new synaptic connections in M1 region of the motor cortex (29). Therefore, we assessed how learning-associated spine formation in deep-layer excitatory neurons (thy1-YFP) varied with GR-PO_4_ using transcranial 2-photon microscopy (Figure 2A). As expected, the majority of spines were stable such that only a small subset of spines was dynamic after training (Figure 2B). Rates of spine formation (Figure 2C) and elimination (Figure 2D) were undistinguishable between WT and KI mice when untrained. But after training KI mice exhibited reduced spine formation (Figure 2C) and excessive spine elimination (Figure 2D) that corresponded with a net decrease of total spine number compared to WT littermates (Figure 2E). Spine maintenance was also diminished in KI mice. This included both training-induced new spines and pre-existing old spines present before training (Figure 2F). In fact, the more training-induced new spines that survived the consolidation period, the better the retention of motor skill performance (Figure 2G).

**Figure 2.**
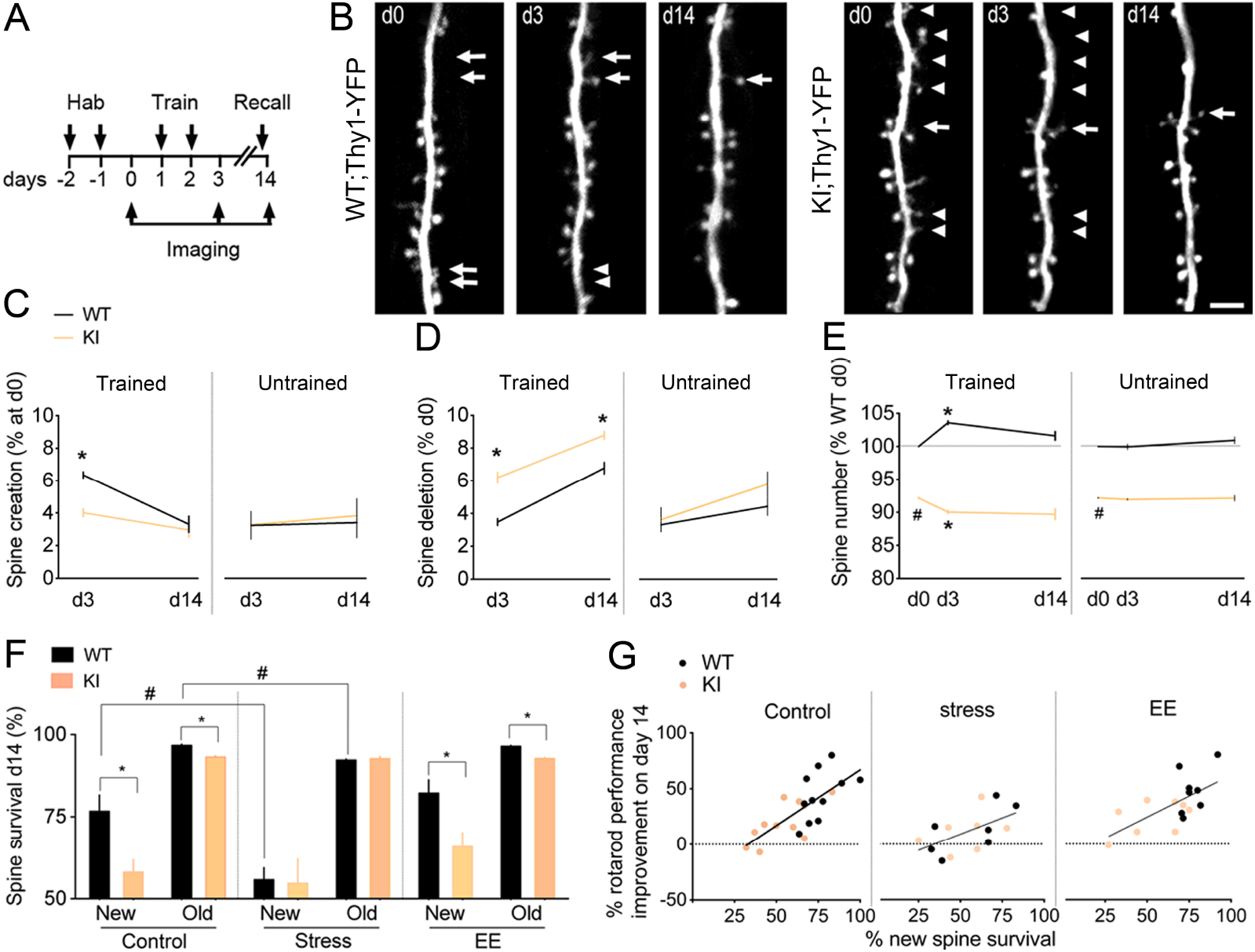
GR-PO_4_ is required for the maintenance of new spines formed at training. (A) Timeline of dendritic spine imaging in M1 cortex of GR-KI;thy1-YFP mice habituated to the non-accelerating rod (2 rpm) for 2 days before training at circadian peak of corticosterone (7 pm) on the accelerating rod (up to 80 rpm). A recall session on the accelerating rod is performed prior to the last imaging session on day 14 (B) Dendritic spines dynamics of L5 principal thy1-YFP neurons in L1 of M1 as a function of genotype and time. Arrows = spine creations; arrowheads = spine deletions. Scale=5 μm. (C) Spine creation in M1 (means ±SEM of n=20 trained WT mice on d0-3, 6 on d14; 18 trained KI on d0-3, 7 on d14 and 6 untrained WT on d0-14, 7 untrained KI on day 0-14). Effect of genotype x training: 3-way ANOVA *F*_1,69_=4.39, *p*<0.05 post-hoc Tukey test WT vs KI \**p*<0.0005. (D) Spine deletion in M1 (means ±SEM of n=20 trained WT mice on d0-3, 6 on d14; 18 trained KI on d0-3, 7 on d14 and 6 untrained WT on d0-14, 7 untrained KI on day 0-14). Effect of genotype: 3-way ANOVA *F*_1,69_=23.28, *p*<0.0001, genotype x training *F*_1,69_=5.3, *p*<0.05 post-hoc Tukey test WT vs KI \**p*<0.05. (E) Spine number in M1 relative to WT control (means ±SEM of n=20 trained WT mice on d0-3, 6 on d14; 18 trained KI on d0-3, 7 on d14 and 6 untrained WT on d0-14, 7 untrained KI on day 0-14). Effect of genotype: 3-way ANOVA *F*_1,116_=2453, *p*<0.0001, genotype x training *F*_1,116_=55.18, *p*<0.0001, post-hoc Tukey test WT vs KI ^#^*p*<0.0001, d0 vs d3 \**p*<0.0001. (F) Survival of training-induced new spines and pre-existing old spines in M1. Means ±SEM of n=11 control mice/ group, 7 stress/ group; 8 EE/ group, 3-way ANOVA: effect of stress *F*_1,64_=8.1, *p*=0.059 post-hoc Tukey test for WT ^#^*p*<0.005. Pairwise comparisons by unpaired *t*-test for the effect of genotype on control new spines *t*_(20)_=2.99 \**p*=0.007 and control old spines *t*_(20)_=6.9 \**p*<0.0001; on new spines EE *t*_(14)_=2.78 \**p*=0.014 and old spines EE *t*_(14)_=7.93 \**p*<0.0001. See Figure S6 for spine addition and elimination data. (G) Correlation between survival at day 14 of spines that formed during training on day 2 and memory retention on day 14 in control conditions (Pearson *r*= 0.73, *p*=0.0001 n=11 WT, 11 KI), chronic stress (*r*= 0.56, *p*=0.036, N=7 WT, 7KI) or EE (*r*= 0.64, *p*=0.0006, n= 8 WT, 8 KI).

To ensure that BDNF-dependent GR-PO_4_ effect on experience-dependent spine plasticity was cell autonomous in excitatory neurons of M1 cortex, we exclusively targeted this set of neurons by *in utero* electroporation (Figure S6A-C). Substitution of endogenous GR with the PO_4_-deficient GR mutant as previously described (30), decreased spine formation and increased spine elimination in the layer 1 of M1 cortex after the training that is consistent with a net decrease of spine density observed in the KI mice. This indicates that the effect of GR-PO_4_ is cell-autonomous.

Chronic unpredictable stress in WT mice decreased spine formation, increased spine elimination (Figure S6D, E) during training, and reduced spine survival during consolidation period, thus mimicking the effect of the KI on spine dynamics, motor skills learning and memory. KI mice showed net spine loss exaggerated with training and no further impact of chronic unpredictable stress on spine formation, elimination and consolidation (Figure 2F, S6F). The lack of any additive effects between chronic stress and deletion of the GR-PO_4_ sites on spine plasticity and indicates a functional redundancy of these pathways. This contrasts with the lack of effects of enrichment living on spine elimination, survival and motor performance (Figure 2F,G). Remarkably, enrichment living reversed spine formation defects in KI mice (Figure S6D, E). However, this did not enhance motor retention because the new spines were unrelated to the training. This is in agreement with a role of BDNF-dependent GR-PO_4_ on the maintenance of training-dependent new spines for better retention of motor performance (Figure 2G).

### BDNF-Val66Met polymorphism recapitulated synaptic and motor defects of GR-PO_4_ deletion

The BDNF-Val66Met polymorphism impairs activity-dependent secretion of BDNF and manifests as defective motor skill training in rodents and humans (28, 31). This phenotype is similar to that observed in the KI mice. Therefore, we tested functional epistasis between KI and BDNF-Val66Met to determine whether both pathways converge during motor skills learning. We found that BDNF-dependent GR-PO_4_ was reduced in mouse carriers of the BDNF-Val66Met allele (Figure 3A). We then crossed heterozygous KI;thy1-YFP mice with heterozygous BDNF-Val66Met;thy1-YFP mice to obtain homozygotes. Four groups of mice: 1) WT, 2) KI, 3) M (BDNF-Val66Met) and 4) M;KI (KI::BDNF-Val66Met) were subjected to *in vivo* imaging of dendritic spines upon motor skill training. M mice exhibited normal acquisition but impaired retention of the motor task. Importantly, the M;KI double mutant mice showed no additional effect (Figure 3B). As expected, spine plasticity in the M mice was defective during training but was unaffected in the untrained groups (Figure 3C). Defects in spine elimination and spine survival in the M mice recapitulated that of the KI mice, with no additive effects in the M;KI double mutants (Figure 3C,D). Together, these results indicated that KI mice and M (with impaired activity-dependent BDNF secretion) have a reduction in their ability to stabilize the new training-induced spines and retain new motor skills. This is consistent with the convergence of both pathways towards new spine consolidation.

**Figure 3.**
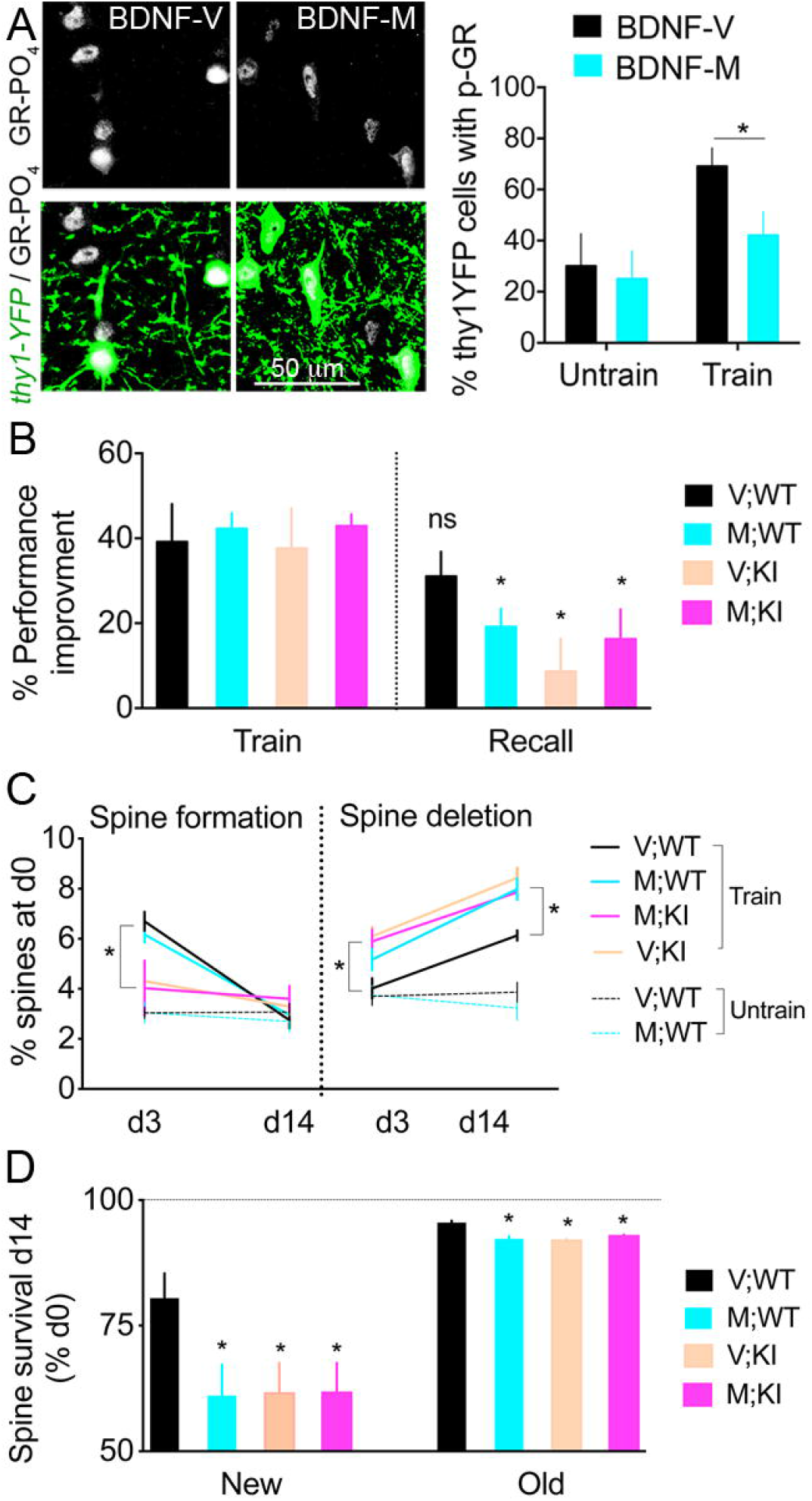
BDNF-dependent GR-PO_4_ is required for training-evoked spine survival and motor skill retention. (A) Effect of BDNF val66met polymorphism on GR-PO_4_ staining in thy1-YFP neurons of M1 cortex after 2 days of rotarod training. Means ±SEM of n=5 untrained mice/group and 7V (BDNF-Val/Val), 8M (BDNF-Met/Met) trained, 2-way ANOVA *F*_1,21_=7.75, ^#^*p*=0.011. Pairwise group comparison between BDNF-M and BDNF-V by unpaired *t*-test *t*_(13)_=2.24 \**p*<0.05. (B) Effect of genotypes on rotarod performance at recall on day 14 expressed as percentage of day 1 in GR-KI mice backcrossed with the BDNF-Val66Met background. Means ±SEM of n=8 V;WT mice, 8 M;WT, 9 V;KI and 11 M;KI. Effect on retention: 3-way ANOVA *F*_1,64_=23.68, *p*<0.0001 post-hoc Tukey test \**p*<0.05. (C) Spine formation in M1 (means ±SEM of n=7 V;WT, 7 M;WT, 7 V;KI and 8 M;KI trained mice and 8 V;WT, 5 M;WT untrained mice). Effect of genotype 2-way ANOVA *F*_3,50_=4.65 and time *F*_1,50_=91.55, *p*<0.01 post-hoc Tukey test \**p*<0.0001. Spine deletion. Effect of genotype: *F*_3,50_= 12.66 and time *F*_1,50_=73.12, *p*<0.0001 post-hoc Tukey test \**p*<0.05. (D) Survival of new spines in M1 formed on day 3 and old spines formed before day 0 (means ±SEM of n=7 V;WT, 8 M;WT, 9 V;KI and 11 M;KI mice). Effect of genotype: 2-way ANOVA *F*_3,62_=2.88, *p*<0.05. Pairwise comparisons by unpaired t-test between V;WT and M;WT on new spines *t*_(13)_=2.28 and old spines *t*_(13)_=4.86; V;WT and V;KI on new spines *t*_(14)_=2.24 and old spines *t*_(14)_=9.95; V;WT and M;KI on new spines *t*_(16)_=2.15 and old spines *t*_(16)_=7.91, \**p*<0.05.

### Poorer retention was consistent with weaker LTP in motor cortex upon GR-PO_4_ deletion

Potentiation of excitatory synaptic transmission in the M1 motor cortex is required to acquire and retain new motor skills (32). To test whether KI mice showed defects in synaptic plasticity, we performed tetanus-induced long-term potentiation (LTP) 1 day post-training. LTP can be induced in WT and KI mice regardless of training (Figure 4A,B). After 3 consecutive tetanus stimulations, LTP saturated in the trained cortex of WT but not KI mice (Figure 4B,C). These data suggested that training occluded LTP in WT but not KI mice (Figure 4D). In fact, LTP occlusion after serial inductions can predict retention of motor skills learning in rodents and humans (33–35). In the KI mice, we find that LTP occlusion was weaker and retention was poorer compared to WT littermates (Figure 4E). This indicates that GR-PO_4_ is essential for functional strengthening of M1 synapses post-training.

**Figure 4.**
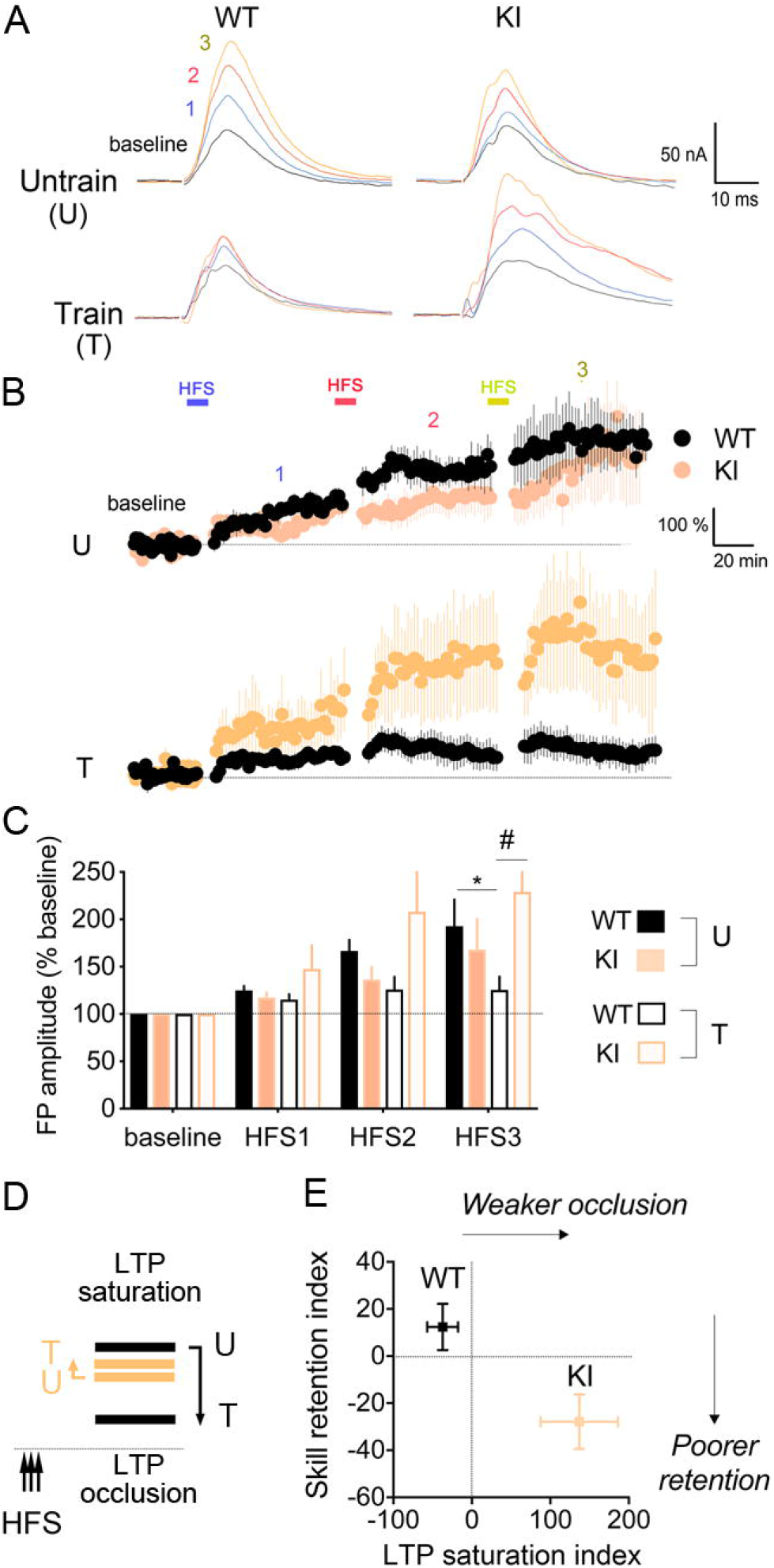
GR-PO_4_ is required for training-evoked plasticity in motor cortex. (A) Field potential (FP) recordings of high-frequency stimulation (HFS)-induced LTP in M1 cortical slices as a function of genotype and training for 2 days on the rotarod. Acute slices on the next day of training are stimulated with half the intensity to reach maximal response in L2 parallel fibers at ≈500 μm distance from the recording electrodes in L2 of M1 cortex. There is a 20 min baseline recording between HFS. (B) FP recordings of tetanus-induced LTP in M1 cortical slices from 8 WT and 5 KI trained mice and 5 WT and 6 KI untrained mice. Means normalized to baseline ±SEM. (C) Averaged FP amplitudes after 3 consecutive HFS (Means of last 40 min/epoch ±SEM). LTP occlusion after HFS3 by unpaired t-test in WT \**p*=0.034. Three-way ANOVA: effect of HFS *F*_3,80_=13.16, *p*<0.0001, genotype *F*_1,80_=4.57, *p*<0.05, genotype x training *F*_1,80_=14.18, *p*<0.005 post-hoc Tukey test ^#^*p*=0.011. (D) Working model: Training in WT but not KI mice occluded LTP saturation. Dashline represents baseline transmission in M1. (E) Weaker LTP occlusion in KI mice corresponded with poorer retention of motor skills. Motor skill retention and LTP saturation indexes are described in the methods section. Effect of genotype on LTP occlusion by unpaired t-test *p*=0.0055 (means±SEM *t*_(14)_=3.28, n=8 mice/ group) and on motor skill retention *p*=0.014 (means±SEM *t*_(22)_=2.64, n=12 mice/ group).

### GR-PO_4_ is required for synaptic mobilization of phosphorylated GluA1

Synaptic delivery of AMPA-type glutamate receptors is thought to contribute to LTP (36). In motor cortex, skill training drives AMPA-type glutamate receptor subunit 1 (GluA1) to postsynapses, contributing to the dynamic changes of glutamatergic synaptic strength (32). Therefore, we tested whether GR-PO_4_ and training have an impact on GluA1 synaptic content and phosphorylation because these regulate AMPA conductance and synaptic strength (37, 38). We first determined if there was a difference in levels of synaptic GluA1 between trained and untrained groups. Synaptosomes were isolated from M1 cortex collected 45 min after training, and synaptic GluA1 levels determined using Western blot analysis (Figure S7). Levels of GluA1 were similar in all groups. However, the level of GluA1 phosphorylation at S831 increased as a function of training in WT but not in KI mice. As GluA1 phosphorylation at this site have been linked to synaptic potentiation in M1 (39), its lack of increase post-training could reflect LTP defects observed in KI mice post-training.

We next investigated cell surface delivery of GluA1 using an established biotinylation assay in cultured cortical neurons (Figure S8A and (40)). Avidin pulldown of biotin-tagged GluA1 newly inserted at the cell surface was detected by Western blot analysis (Figure 5A). Upon co-stimulation of the BDNF and GR pathways, levels of GluA1 and its phosphorylation isoform were lower in neurons expressing GR-KI mutant compared to GR-WT (Figure 5B). This effect was specific of GluA1 (Figure S8B). These results suggest that GR-PO_4_ at the time of training promotes the mobilization of synaptic GluA1. Does this permit the structural maturation and stabilization of training-dependent spines?

**Figure 5.**
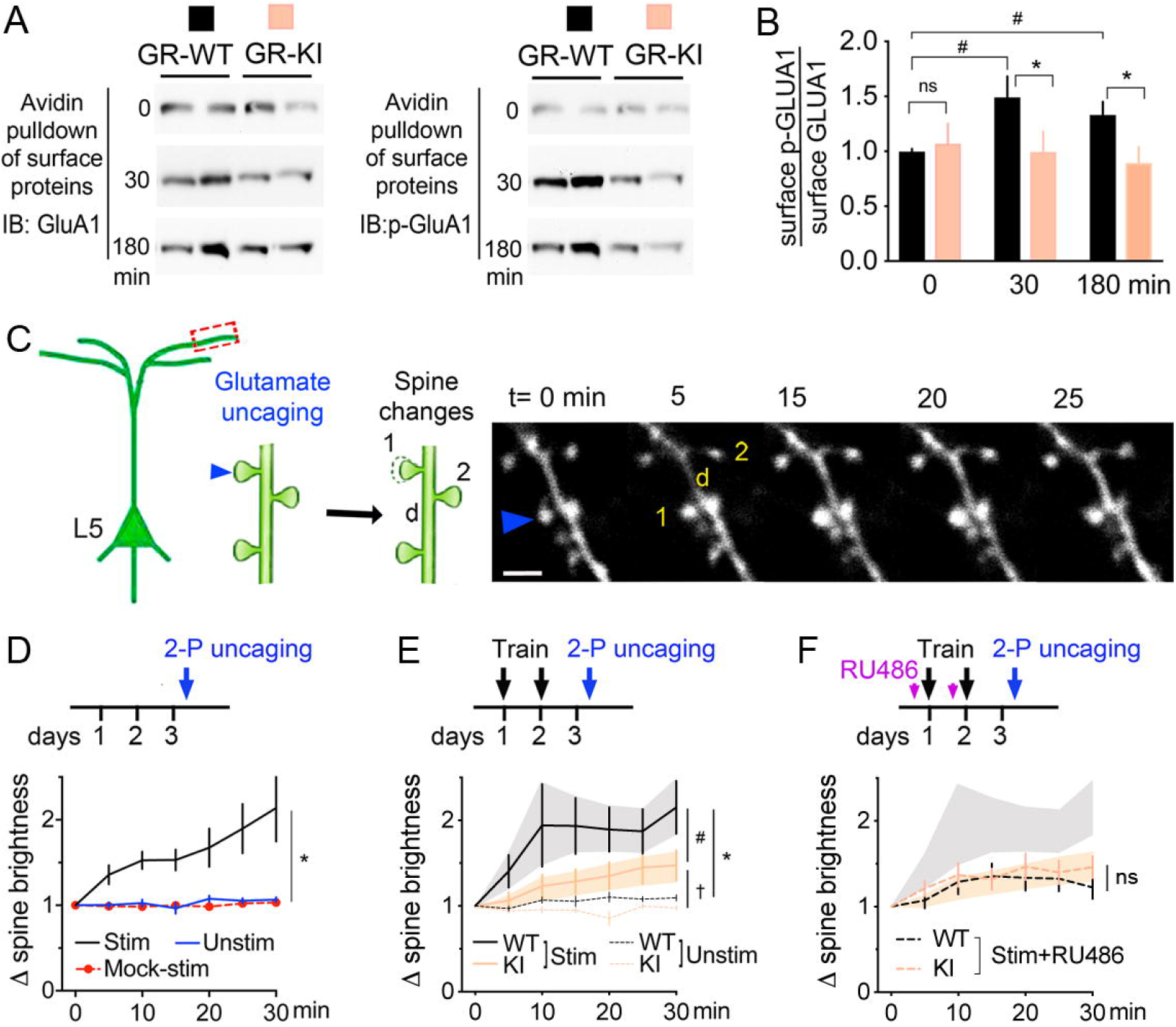
GR-PO_4_ is required for training-evoked spine maturation. (A) Avidin pulldown of biotinylated proteins from primary DIV14 neurons electroporated with GR-WT or GR-KI at DIV0, and stimulated with 25 ng/ml BDNF and 1 μM dexamethasone (Dex) for the indicated time. Each column of immunoblots (IB) for GluA1 and p-GluA1 represents an independent sample. (B) Surface G1uA1-PO_4_/G1uA1 in cortical neurons expressing GR-WT or GR-KI. Means ±SEM of n=11 WT and 8 KI/group, 2-way ANOVA: Effect of mutant *F*_1,53_=6.61, *p*=0.016. Pairwise group comparison by unpaired t-test \**p*<0.05, ^#^*p*<0.001. (C) *In vivo* 2-photon uncaging of 20 mM MNI-glutamate (90 iterations laser at 720 nm, 0.7 mW, 0.1Hz) in HEPES-buffer in M1. Arrow indicates photostimulated spine (1) compared to underlying dendritic shaft (d) and neighboring unstimulated spine (2). Scale=3μm. (D) Specificity of glutamate-evoked spine enlargement normalized to baseline spine size in WT mice. Means ±SEM of n=10 stimulated spines, 9 mock-stimulated spines, 20 neighbor spines in 3 mice, 2-way ANOVA: Effect of stimulation *F*_2,237_=83.9, *p*<0.0001; time *F*_6,237_=6.78, *p*<0.0001 post-hoc Tukev test \**p*<0.0001. (E) Effect of genotype on glutamate-evoked spine enlargement after 2 days of rotarod training. Means ±SEM of n=23 stimulated spines WT, 40 stimulated spines KI, and 18 neighbor spines WT, 20 neighbor spines KI in 4 mice/ group, 2-way ANOVA: Effect of stimulation *F*_3,631_=21.93, *p*<0.0001; time *F*_6,631_=2.43, *p*=0.024 post-hoc Tukey test ^#^*p*<0.0001, \**p*<0.0001, ^†^*p*=0.009. (F) Effect of RU486 (20 mg/kg, IP 20 min before each training session) on glutamate-evoked spine enlargement. Means ±SEM of n=17 stim+RU486 WT spines, 13 stim+RU486 KI spines in 3 mice, 2-way ANOVA: Effect of stimulation *F*_3,605_=10.13, *p*<0.0001; time *F*_6,605_=3.96, *p*=0.0007.

### GR-PO_4_ pathway is required for strengthening glutamatergic response post-training

Training-induced LTP occlusion has been previously associated with enlargement of dendritic spine heads in upper layers of motor cortex, a process dependent on glutamate (41). Therefore, we tested the role of GR-PO_4_ on glutamate-dependent structural maturation of dendritic spines *in vivo*. To this end, we used time-lapse imaging of dendritic spines and 2-photon uncaging of MNI-glutamate directly into M1 cortex through a cranial window (Figure 5C). Glutamate uncaging on spine head causes its enlargement without affecting unstimulated neighboring spines (Figure 5D). Photo-stimulation of spines in absence of MNI-glutamate had no effect, confirming the specificity of glutamate uncaging response. In trained mice, the rate of glutamate-evoked spine enlargement was 79 ± 11% in WT and 59 ± 8% in KI. Glutamate-evoked spine enlargement is also slower in KI mice than in WT mice (Figure 5E). Thus, glutamate-induced spine maturation is weaker in KI mice than WT mice.

Pharmacological blockade of GR with RU486 administered immediately before training impaired the glutamate response of dendritic spines in WT controls, whereas in KI mice no further inhibition of the response was observed (Figure 5F). The lack of additive effects between the deletion of GR-PO_4_ and GR inhibition indicated functional redundancy. In contrast, RU486 administered after training had no effect on the glutamate response of dendritic spines (Figure S9). This further indicates that BDNF-dependent GR-PO_4_ at training is required for the strengthening of the glutamatergic response of dendritic spines after training.

## DISCUSSION

Here we provide the first *in vivo* evidence for the dependence of GR-PO_4_ on BDNF activity-dependent release for remodeling excitatory synapses in the cortex upon learning. Our study reveals that either the lack of GR-PO_4_ induced by mutation of the BDNF-dependent phosphorylation sites, or decreased activity-dependent BDNF secretion modeled in the BDNF-Val66Met mutant reduced the survival of learning-associated new spines and impaired memory retention. Moreover, combining the BDNF-Val66Met with the GR-PO_4_ mutant mice does not induce further deficits on spine maintenance and behavior as the single mutants, indicating that the effect of GR-PO_4_ on spine dynamics occurs via BDNF. In agreement with this finding is an increase in GR-PO_4_ in the motor cortex by motor skills training, and reduced sensitivity to corticosterone and stress in humans and mice with the BDNF-Val66Met allele (20, 42).

Our findings also shed light onto the molecular mechanisms underlying how information in the brain is stored (i.e. motor engram) in response to external stimuli including stress and enriched environment. We observed that chronic stress increased the turnover of spines in wild type but not in the GR-PO_4_ deletion mice as both stress and GR-PO_4_ deletion resulted a net loss of training-induced new spines in the cortex. This highlights the critical role of BDNF-dependent GR-PO_4_ on the maintenance and connectivity of new training-induced spines within a motor engram. Our findings also predict that physical exercise and/or an enriched environment would promote spine maintenance in cortex by transforming the training-related new spines into a persistent pool of memory spines, and alleviate the impact of stress on the synaptic engram (43, 44). Motor learning was also associated with an increase in the active phosphorylated form of the GluA1 into synapses that stabilize and strengthen training-related newly formed spines and motor skill retention (37, 45). These effects were reduced in GR-PO_4_ deletion mice and recapitulated by GR inhibition with RU486 or chronic stress, which diminished GR-PO_4_ (46–48). Likewise, AMPAR inhibition with CNQX injection in M1 cortex reduced motor skill retention (32), suggesting that maintenance of the synaptic memory engram depends on GR-PO_4_ signaling to AMPAR (25, 49). Consistent with the effect of GR-PO_4_ deletion, a stable deficit of motor skill retention was also found after ablation of Drd1-neurons in dorsolateral striatum (targets of L5 pyramidal neurons) during consolidation (50). D1/D5 antagonist or dopamine depletion in motor cortex increased spine elimination and decreased spine survival in motor cortex such as in the KI mice (51). In fact, D5 controls AMPA currents in motor cortex by promoting GluA1 phosphorylation such as GR-PO_4_ (52, 53). The similarities with the effect of GR-PO_4_ deletion suggest a putative link between GR-PO_4_ and dopamine signaling that will need further studies.

Physiological consequences of the BDNF-Val66Met allele, like that of GR-PO_4_ deletion, are revealed not in the basal state but in response to experience-driven increase of neural activity (28, 54). This indicates that GR senses and responds to BDNF by selective GR phosphorylation that alters its activity and promotes learning and memory. Whether this reflects changes in GR genomic and/or non-genomic activity remains to be determined. Evidence obtained from RU486 injections indicate that GR-PO_4_ signaling is required hours rather than minutes before dendritic spine mature via a glutamate dependent mechanism. Therefore, these effects depend on a slow process, likely genomic to consolidate new spines. This could have implications for the encoding and the persistence of new structural engrams required for adaptive plasticity that either translate to a new behavior or adapt to a changing environment (i.e. stress, learning, enrichment).

Our findings have also translational implications. Imbalances in the integration of BDNF-GR signaling axes would be predicted to trigger maladaptive processes that contribute to the pathophysiology observed in many neurological and other diseases (18, 24, 55, 56). For example, glucocorticoid receptor insufficiency is associated with a state of low BDNF levels in diseases presenting an activated inflammatory system that is relevant for patients with major depression, schizophrenia, glucocorticoid resistance syndrome, Alzheimer’s disease (57–60). Antidepressant therapies that successfully increase BDNF levels also correct GR-PO_4_ and glucocorticoid receptor sensitivity (23, 61). GR-PO_4_ has been suggested to be a mechanism contributing to glucocorticoid resistance in multiple disease models (62). Although a majority of GR-PO_4_ sites is glucocorticoid-dependent (63), our data indicate that GR-PO_4_ can also be glucocorticoid-independent. This implies that GR activity is influenced by contextual signals in addition to glucocorticoids (22, 26, 64). This applies, for example, to the maintenance of sensory motor engrams as long as activity-dependent BDNF secretion and glucocorticoid oscillatory pulses are synchronized and stimulate the GR-PO_4_ pathway. For that reason, aligning glucocorticoid treatment to neurotrophin release should be considered to promote neuroplasticity for sensory motor rehabilitation.

## METHODS

All experiments were carried out in accordance with the Directive by the Council of the European Communities (86/609/EEC) and approved protocol (00651.01) following institutional guideline for the care and use of laboratory animals. All tools are listed in table S1.

### ANIMALS

PO_4_-delicient GR mouse (Flex GR-A152/A284 knockin by Ozgene Pty Ltd) was generated in C57Bl6 background and crossed with constitutive ROSA26-FLP line (*Gt(ROSA)26Sor^tm2(FLP*)Sor^*) to remove the neomycin cassette and constitutive ROSA26-CRE line (*Gt(ROSA)26Sor^tm1Sor^*) to produce a general deletion of GR-PO_4_ sites (for details, see Figure S1). Thy1-YFP transgenic mice (*B6.Cg-Tg(Thy1-YFP)HJrs/J*), BDNF Val66Met mice (*Bdnf^tm1Flee^*, MGI:3664862) and wildtype mice in C57BL6 background (Jackson labs) were housed under a 12 hr light/ dark cycle (on 6 am, off 6 pm) with unrestricted access to food and water. Chronic unpredictable mild stress includes one of the following daily random stressors (wet bedding, no bedding, food deprivation, crowded cage, 2h or 6h restrain, tilted cage, shaking, 24-hr light cycle, forced swim, tail suspension) for 10 consecutive days. Enrichment consisted of larger cages with toys of various colors and textures (tubes, tunnels, ladders, Lego^™^, beads, nest) for 10 consecutive days. Homozygous males produced by heterozygous breeding schemes were used in all protocols. All efforts were made to minimize animal suffering and to reduce the number of mice utilized in each experiments.

### OPEN FIELD

One month old mice freely explored an arena (50□cm × 50□cm) for 10□min. Movement paths were captured by an infrared digital camera and total distance travelled determined with Ethovison XT software (Noldus). Thigmotaxis represents the time spent in the center of the arena (29 cm × 29 cm).

### ELEVATED PLUS MAZE

Mice at 3 month of age freely explored the arms (elevation 50 cm × 50 cm × 20 cm) for 10 min. Movement paths were captured with a digital camera to score the number of entries and time spent in each arm.

### FORCED SWIM

Mice at 3-months of age were subjected to forced swim test (FST) for 9 min in a beaker (15 cm × 25 cm) filled with tap water at room temperature during the stress protocol (once a week between postnatal day PND 23 and 36) and video recorded trial. Trial data represent an acquired behavioral response in the stress groups but a novel response in the control groups reared in standard living conditions.

### TAIL SUSPENSION

Mice at 3-months of age were subjected to tail suspension test (TST) for 6 min during the stress protocol (once a week between PND 23 and 36) and video recorded trial. Trial data represent an acquired behavioral response in the stress groups but a novel response in the control groups reared in standard living conditions (65).

### MOTOR LEARNING

Mice were habituated for 30 minutes (15 trials) on the non-accelerating rotarod (2 rpm, 1 min followed by 30 sec rest inter trials) for 2 consecutive days at glucocorticoid circadian oscillation peak (7 pm in darkness) before 2 training sessions of 1 hour also at 7 pm in darkness. Training consisted in 15 trial sessions on the accelerated rod (from 2 to 80 rpm reached in 2 min with 1 min rest inter trial) for 2 consecutive days. Control mice are habituated and trained on a slow accelerating rod (2 rpm) that did induce spine patterning in the motor cortex. To characterize training-induced c-FOS expression, mice were euthanized 45 min after training at 80 rpm (trained group) or 2 rpm (untrained group). Motor performance is indexed as latency to fall. Acquisition is calculated as performance improvement index = mean d_2_(t_14_+t_15_) – mean d_1_(t_1_+t_2_). Retention on recall (7 pm) is calculated as motor skill retention index = improvement index – (mean d_14_(t_1_+t_2_) – mean d_1_(t_1_+t_2_)) where d is the day at testing and t is the trial number.

### TRANSCRANIAL 2-PHOTON MICROSCOPY

Mice were anesthetized with a mix of ketamine/ xylazine (0,075mg/g and 0,01mg/g, respectively) prior to surgery. Thin-skull preparation for transcranial *in vivo* imaging in motor cortex (coordinates from Bregma −1.3 mm, +1.2 mm lateral) was preferred to open skull preparation because it prevents artifacts due to surgical-induced chronic inflammation (66, 67). All images were acquired with a Zeiss LSM710 2-photon microscope coupled to a Ti:sapphire laser tuned to 920 nm (Spectra physics) and a water immersion 20x objective (NA 1.0, Apocromat, Carl Zeiss). Laser power was kept below 5 mW to avoid photodamage. Images were taken at each image session for each mouse using 70 μm × 70μm (512 × 512 pixels) at 0.75 μm step with a scanning dwell time of 2.55 μsec/ pixel.

### RE-IMAGING OF THE SAME FIELD OF VIEW

Repeat imaging of the same dendritic field before (image 1 at PND23) and after training (image 2 at PND26), as well as at recall (image 3 at PND36) offers an opportunity to identify a synaptic engram of motor skill training that correlates with motor performance (25). Comparison of pairs of images identified stable spines (present in image 1, 2 and 3), eliminated spines (present in image 1 but not in image 2 and/or 3), formed spines (present in image 2 but not in image 1) and survival of new spines (present in image 2 and 3 but not in image 1) as well as pre-existing old spines (present in image 1 but not in image 2 or 3). To this end, a detailed map of the pial vasculature was taken for subsequent relocation. Bone regrowth between imaging session is thinned using disposable ophtalmic surgical blades (Surgistar). Skull is further thinned between imaging sessions down to 18-20 μm allowing no more than 4 consecutive sessions to avoid cracking the skull (67). Scalp is sutured and topped with topical antibiotic cream.

### TIMELASPE IMAGING THROUGH A CRANIAL WINDOW

Two-photon microscopy was performed at open skull preparations to facilitate access of drugs to the pial surface of cortex after surgical removal of meninges covered by a thin layer of low-melting point agarose (1.5% in HEPES-buffered artificial cerebrospinal fluid (ACSF) to avoid heartbeat motion artifacts. Baseline spine dynamics were captured in HEPES-buffered ACSF (in mM: 120 NaCl, 3.5 KCl, 0.4 KH2PO4, 15 glucose, 1.2 CaCl2, 5 NaHCO3, 1.2 Na2SO4, HEPES, pH=7.4) for 5 min prior glutamate uncaging was performed in HEPES-buffered ACSF with 20 mM MNI-glutamate (Sigma) photostimulated with 90 iteration at 0.1Hz of 5 % laser power tuned to 720 nm (0.7 mW) as described (68). Success rate of spine enlargement was calculated as the number of trials in which spine brightness exceeded 10% of initial value divided by the total number of trials.

### IMAGE ANALYSIS

All images obtained from time lapses imaging sessions were realigned with Image J plugin RegStack to minimize artifacts of heartbeat pulsations. Images were stitched together using ImageJ. Image stacks in between sessions were compared using ImageJ. Dendritic segments included in the analyses met the following criteria: (1) be parallel or at acute angles relative to the coronal surface of sections to allow unambiguous identification of spines, (2) segments had no overlap with other branches, (3) dendritic segments from apical tree were imaged within the first 100 μm from the pial surface. About 200 dendritic spines from at least 20-30 dendritic segments were counted per conditions throughout the imaging sessions and averaged per animal. For this study, more than 23,000 spines were tracked in time-lapse images (3 to 5 time points). The resolution of our 3D images is insufficient to resolve spines reliably in the z-axis. So, spines below or above dendrites were not analyzed. All clear protrusions emanating laterally from the dendritic shaft, irrespective of apparent shape, were counted. Spines were considered deleted if they disappeared into the haze of the dendrites (length < 5 pixels); spines were considered formed if they clearly protruded from the dendrites (length ≥ 5 pixels). To calculate spine brightness (the pixel values containing the spine head were summed. Background fluorescence, calculated over the same-sized box adjacent to the spine, was subtracted. Since dendritic shaft diameters were constant and relatively uniform, we used them to correct fluorescence levels. Average shaft pixel intensity was calculated over the same-sized box adjacent to the spine. The background-subtracted, pixel value for each spine was divided by the average shaft pixel value as previously described (69). The resulting relative brightness is expected to be proportional to the spine volume. Net ratios, the fraction of spines gained subtracted of the spines lost day to day, were calculated as NR = (N_formed_ - N_deleted_)/(2 × N_total_).

### LIVE SLICE PREPARATION

Mice were decapited at PDN26, 12-15hrs after the last training session as described (39) and brains immersed in ice-cold oxygenated (95% O_2_, 5% CO_2_) ACSF containing (in mM): 127,25 NaCl, 1,75 KCl, 1.25 KH_2_PO_4_, 1 MgCl_2_, 2 CaCl_2_, 26 NaHCO_3_, 10 glucose. Coronal slices (400 μm) including the M1 area (1.5–3.5 mm anterior to Bregma, 2–4 mm lateral) were transferred to a temperature controlled (34 +/-1□ C) interface chamber and perfused with oxygenated ACSF at a rate of 1mL/min. Slices were allowed to recover for at least one hour before recordings.

### ELECTROPHYSIOLOGICAL RECORDINGS

Stimulation electrodes were positioned in L2/3, 1.5-1.8 mm lateral to the midline, and recording electrodes placed 500 μm laterally. Field potentials (FP) were evoked by stimulation of 0.2 ms at 0.03 Hz. Induction of long-term potentiation (LTP): stimulus intensity eliciting 50% of the maximum amplitude was used for all measurements before and after LTP induction paired with a touch application of bicuculline methiodide (3.5 mM, GABA_A_ antagonist) as described (33). Baseline amplitudes were recorded using single stimuli applied every 30 sec. Following a 30-min stable baseline period, LTP was induced by theta burst stimulation (TBS), consisting of 10 trains of 5 Hz stimuli, each composed of 4 (200 msec) pulses at 100 Hz, repeated 5 times every 10 sec. Maximum LTP values are expressed as percentage of baseline. Neural pathways were considered saturated if the difference between two states of LTP inductions in the habituated and trained mice were not significantly different (*P*>0.5). The magnitude of occlusion of LTP-like plasticity was calculated as occlusion index = (d_2_(LTP_1_-baseline) - mean d_0_(LTP_1_-baseline)) + (d_2_(LTP_2_-LTP_1_) - mean d_0_(LTP_2_-LTP_1_)) + (d_2_LTP_3_-LTP_2_) - mean d_0_(LTP_3_-LTP_2_)) where d_0_ is after 2 days of habituation (2 rpm) and d_2_ is after 2 days of training (80 rpm).

### IMMUNOHISTOCHEMISTRY

Mice were anesthetized at PDN26 or PDN36 with pentobarbital (50 mg/kg, i.p., Ceva santé Animale) and perfused at a rate of 3 ml/min through the ascending aorta with 30 ml of ice-cold 0.9% NaCl prior to decapitation. Brain hemisections were fixed with 4 % ice-cold paraformaldehyde for 2 h and sectioned with a vibratome. Free-floating coronal sections rinsed in PBS were blocked in 3 % normal donkey serum, PBS, 0.1 % triton X-100 for 2 h at 25 °C. Antibodies are listed in table S1. Images were acquired with a confocal microscope LSM780 (Carl Zeiss) and 10x, 20x dry objectives and 40x oil-immersion objective to capture dendrites spine for counting densities. Excitation and acquisition parameters were unchanged during the acquisition of all images. More than 26,000 NG neurons, 7,000 PV neurons, 24,000 GR cells and 2,200 thy1-YFP neurons were counted in all groups to determine proportions of cells co-labeled with c-FOS and GR-PO_4_.

### SYNAPTOSOME PREPARATION

Mice were anesthetized at PDN26 with pentobarbital (50 mg/kg, i.p., Ceva santé Animale) and perfused at a rate of 3 ml/min through the ascending aorta with 30 ml of 0.9 % NaCl prior to decapitation. Motor cortex from brain hemisections were harvested from 200 □ m-thick sections dissected with a tissue punch (Stoelting Co.). Potter was used to make homogenates in ice-cold 0.32 M sucrose, 1 mM EDTA, 10 mM HEPES pH=8, 1 mg/ml BSA complemented with protease and phosphatase inhibitors, and centrifuged twice (1,000 g for 1 min) to clear debris. Particles were centrifuged (14,000 g 12 min) and pellet suspended in ice-cold 45 % Percoll diluted in 140 mM NaCl, 5 mM KCl, 25 mM HEPES pH = 8.0, 1 mM EDTA, 10 mM glucose plus inhibitors. Synaptosome fractions was collected at the surface after centrifugation (14,000 g 2 min), rinsed in ice-cold 0.32 M sucrose, 1 mM EDTA, 10 mM HEPES pH=8. After centrifugation (14,000 g 12 min) synaptosomes were lysed in 2% SDS.

### DNA TRANSFECTION

*In utero* electroporations (30 v, pON 50 ms, pOFF 950 ms, five pulses with NEPA21, (Nepagene), 1 μg DNA) were performed at embryonic day 15 on mouse embryos and newborns developed for 1 month of age (23). Mice were anesthesized with 4% isoflurane/oxygen and maintained at 1.5–2% isoflurane (Abbott laboratories) throughout surgery using TEC3N (Anesteo). Mice received preemptive analgesia with Lidocaïne (Xylovet^®^, 3.5 mg/kg at incision site). A subcutaneous injection of the analgesic buprenorphine (Buprecare^®^, 0.05 mg/kg) was administered post-surgery and the next day. *In vitro* electroporations were performed with the AMAXA system. Plasmids electroporated consist of DNA vectors for molecular replacement of endogenous GR by the shRNA resistant PO_4_-deficient GR (GR-2A: S152A/S284A) or GR-WT as previously described (30).

### CELL CULTURE AND BIOTINYLATION ASSAY

Primary E18 cortical neurons prepared from time-pregnant C57BL6 mice, were cultured on poly-D-lysine, and maintained for 2 weeks in vitro in Neurobasal medium containing B27 supplement, 0.5 mM L-glutamine, 10 μM 5-fluorouridine, 10 μM uridine. Cells were rinsed three times with PBS containing 1 mM CaCl_2_ and 0.5 mM MgCl_2_ and incubated successively at 4□ C with sulfo-NHS-acetate (Pierce) to block surface proteins, with 100 mM glycine to quench reaction, at 37□ C for the indicated time in presence of 25 ng/ml BDNF and 1 M dexamethasone, and finally at 4□C with sulfo-NHS-LC-biotin (Pierce) to label newly inserted surface proteins. Cells were lyzed in 10 mM Tris-HCL, pH=8, 150 mM NaCl, 1 mM EDTA, 10% glycerol, 1% NP40, 0.1% SDS plus protease inhibitors and cleared by centrifugation (12,000 g for 10 min). Streptavidin-agarose pulldown (Pierce) permitted purification of biotinylated surface proteins further rinsed five times with RIPA buffer and resolved in 10% SDS/PAGE.

### WESTERN BLOT

Protein concentrations were measured with BCA assay against BSA standards (ThermoFisher Scientific) using a plate reader for measuring absorbance (Tecan). Immunoreactivities (antibodies listed in Table S1) were revealed by chemiluminescence (GE Healthcare) and densitometric analysis of grayscale digital images using Chemidoc Touch (Bio-Rad Laboratories).

### STATISTIC

Parameters used to quantify imaging data include (i) formation/deletion/survival of dendritic spines, (ii) density of dendritic spines, (iii) spine head diameter, (iv) double-labeled cells, (v) cortical layers visualized with DAPI stain and thy1-YEP in L5. Parameters used to quantify motor performance data include (i) latency to fall the rotarod within-trials improvement as index of learning, and (ii) inter-trials improvement as index of retention. Representation of N for each data set is indicated in figure legends. All data collected in animals were from littermate controls and averaged per experimental groups. We used Student’s *t*-test to compare 2 groups or time points, Pearson correlation for linear associations between datasets with Prism 8.0 Software (GraphPad). We used factorial ANOVA to compare multiple groups (training, genotype, stress and enrichment) followed by *post-hoc* pairwise comparison with Tukey test for corrections. All data are shown as means ± standard error of the mean. Significance level is set at α≤0.05. No data were removed from analyses including statistical outliers. Estimates of sample size were calculated by power analysis based on preliminary data. Sample size was chosen to ensure 80 % power to detect the pre-specified effect size. Pre-established criteria for stopping data collection included: (i) mice reaching ethical endpoint limits; (ii) unexpected mortality (e.g. anesthesia); (iii) crack of the thin skull preparation that would cause unwanted inflammation; (v) brains badly perfused and unusable for histology.

## ACKNOWLEDGEMENTS

We thank F.S Lee (Weill Cornell Med School, USA) for providing the Val66Met mice, WB Gan (NYU School of Medicine, USA), C Liston (Weill Cornell Med School, USA) and C Lafont (IPAM) for their advice on 2-photon microscopy. This work is supported by INSERM/AVENIR (FJ), FP7 Marie Curie (FJ), Montpellier Univ (FJ), Fondation pour la recherche médicale (FJ) and the NIH R56MH115281 (MJG, FJ).

## CONTRIBUTIONS

M.A-L and F.J designed methods. M.A-L and Y.D carried imaging studies. M.M produced synaptosomes, Western and histology. A.B and M.D performed LTP recordings. M.J.G produced GR antibodies. M.J.G and F.J designed the KI mouse. M.A-L, M.J.G, F.J wrote the manuscript. All authors approved the final manuscript and declare no competing interests.

## Abbreviations

BDNF: Brain-derived neurotrophic factor
GR: glucocorticoid receptor
LTP: longterm potentiation
PO_4_: phosphorylation
MNI-glutamate: 4-Methoxy-7-nitroindolinyl-caged-L-glutamate
GluAl: glutamate receptor subunit A1

