## supplemental figures for "Persistence of learning-induced synapses depends on neurotrophic-priming of glucocorticoid receptors"

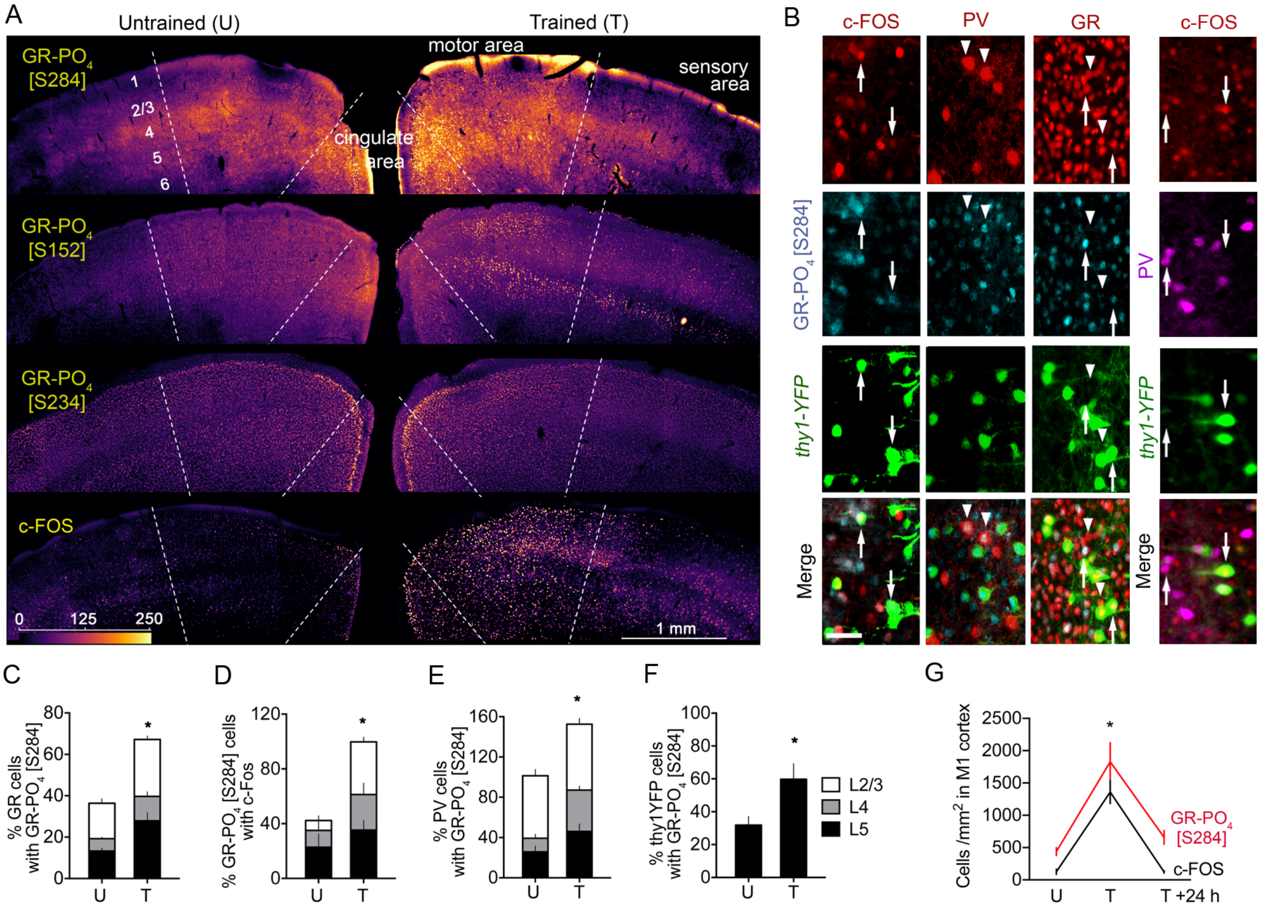


**Figure S1. GR-PO$_4$ induction in neurons of motor cortex upon motor skills training on the rotarod.**

(A) Expression heat map of GR-PO$_4$ isoforms (p-Ser284, p-Ser-152, p-Ser234) and c-FOS in motor areas of cortex 45 min after 2 days of training (T) or no training (U). Induction of GR-PO$_4$ (p-Ser284 and p-Ser-152) and c-FOS are most significant in the layers L2/3 and L5 of M1 cortex.

(B) Cell markers co-expressing GR-PO$_4$ and c-FOS in L5 of M1 cortex. Arrows indicate co-expression of markers. Motor training induced GR-PO$_4$ signaling in principal neurons as showed by triple labeling of thy1-YFP, GR-PO$_4$ and c-FOS in L5 M1 cortex. Motor training also induced GR-PO$_4$ and c-FOS in parvalbumin (PV) interneurons as showed by double labeling of PV, GR-PO$_4$ as well as PV and c-Fos in L5 M1 cortex. Scale=25 µm.

(C) Training induced GR-PO$_4$ in M1 cortex. Proportion of GR-PO$_4$ cells co-labeled with GR. Means ±SEM of n=7 mice/ group, 2-way ANOVA: Effect of training $F_{1,36}$=37.67, $p$<0.0001, post-hoc Sidak test *$p$<0.01.

(D) Training induced GR-PO$_4$ in cells that also express c-FOS in M1 cortex. Proportion of GR-PO$_4$ cells co-labeled with c-FOS. Means ±SEM of n=7 mice/ group, 2-way ANOVA: Effect of training $F_{1,36}$=12.34, $p$=0.0012, post-hoc Sidak test *$p$=0.0065.

(E) Proportion of PV neurons co-labeled with $GR\text{-}PO_4$. Means ±SEM of n=7 mice/ group, 2-way ANOVA: Effect of training $F_{1,36}=13.7$, $p=0.0007$, post-hoc Sidak test $*p<0.05$. Training induced $GR\text{-}PO_4$ in a substantial number of PV cells in M1 cortex.
(F) Proportion of thy1-YFP neurons co-labeled with $GR\text{-}PO_4$. Means ±SEM of n=7 mice/ group. Unpaired t-test: Effect of training $t_{(12)}=2.59$ $*p=0.023$. Training induced $GR\text{-}PO_4$ in a substantial number of principal thy1-YFP neurons in M1 cortex.
(G) Induction of $GR\text{-}PO_4$ and c-FOS 45 min after the last training on day 2 (T, n=5) and 24 hours post-training on day 3 (T+24h, n=4) compared to untrained mice sacrificed on day 2 (U, n=5). Means ±SD, 1-way ANOVA: Effect of training $F_{(5,22)}=18.8$, $p<0.0001$ post-hoc Sidak test $*p<0.001$.

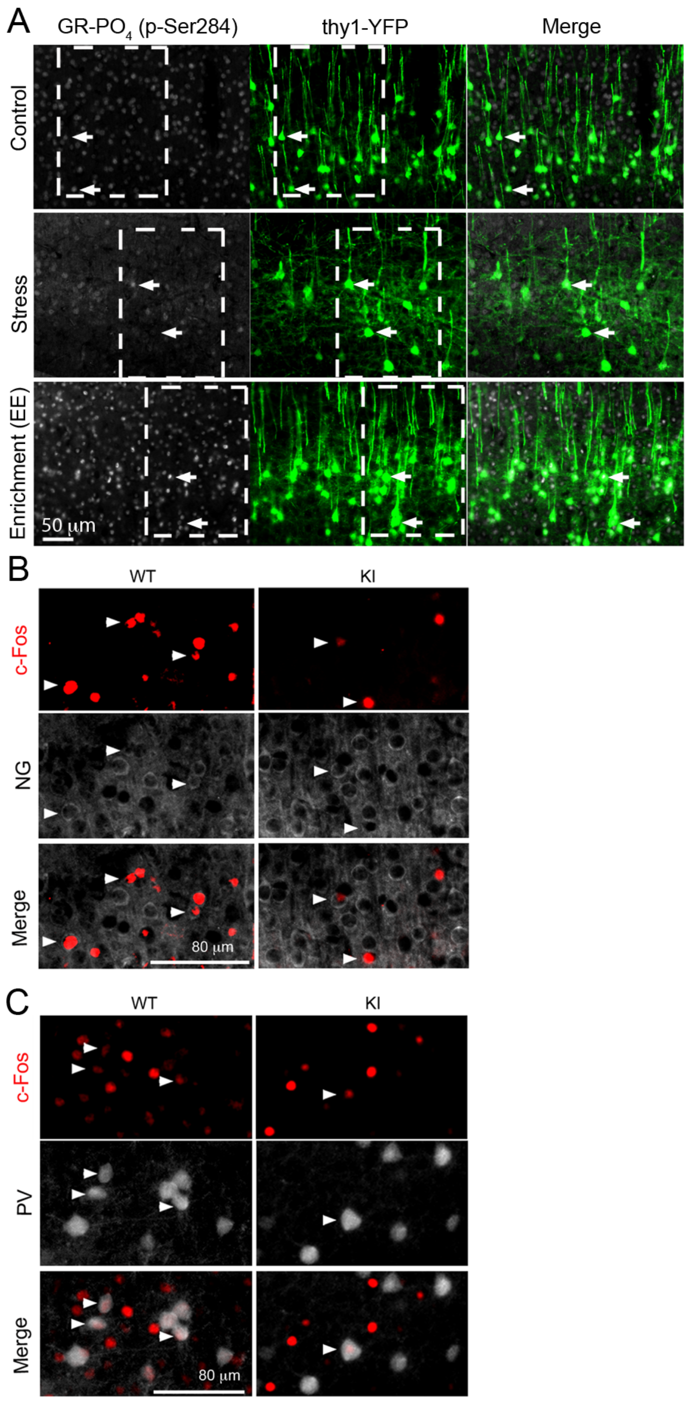


**Figure S2. Induction of GR-$PO_4$ in motor cortex and effects of its deletion.**

(A) Effect of stress and enrichment (EE) between postnatal day (PND)25 and PND36 immediately after the training on GR-$PO_4$ (p-Ser284) in the layer 5 in M1 cortex. Insets indicate the fields of view displayed in figure 1F.

(B) Induction of c-FOS in principal neurons marked with neurogranin (NG) in the layer 2/3 of the M1 cortex from WT and KI mice sacrificed 45 min after 2 days of training (15 trials each). This refers to Figure 1G.

(C) Induction of c-FOS in inhibitory neurons marked with parvalbumin (PV) in the layer 5 of the M1 cortex from WT and KI mice sacrificed 45 min after 2 days of training (15 trials each). This refers to Figure 1G.

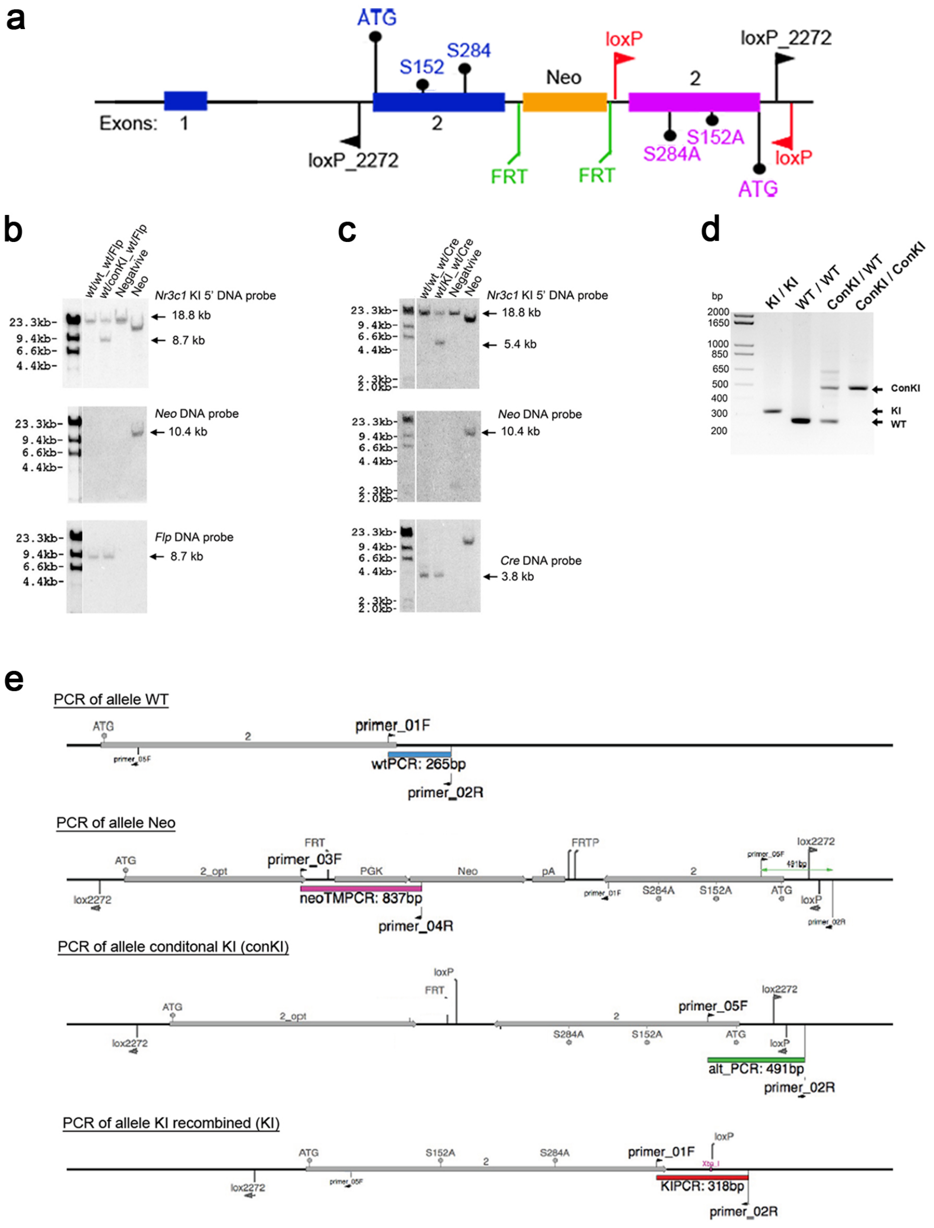


**Figure S3. Knockin construct of $PO_4$-deficient GR mouse at sites responding to BDNF.**

(a) N*r3c1* locus on chromosome 18 (chr18:39410545-39491301) (MGI:95824) presenting an exon 2 with S152 and S284 (WT) and another exon 2 with the mutations A152A and A284 (KI) in head-to-head orientation flanked by 2 pairs of loxP sites, both targets of the CRE recombinase. To avoid the formation of RNA duplex between both exon 2, the nucleic acid sequence of exon 2 WT has been engineered to deviate from the original (mus musculus C57Bl6) while maintaining the aminoacid sequence; the nucleic acid sequence of the exon 2 KI is the original with the exception of S152A and S284A mutations. The neomycin cassette served for the selection of ES clones (derived from C57Bl6 background) that integrated the construct. Selected clones were implanted in C57Bl6 recipient females. F1 were bred with C57Bl6 mice for transmission of the conditional allele (conKI) to the germline. Subsequent cross with constitutive FLP mouse line (ROSA26-FLP by Ozgene) permitted general deletion of the neo cassette. Subsequent cross with constitutive CRE mouse line (ROSA26-CRE by Ozgene) permitted general replacement of WT exon by KI exon. Heterozygous F1 were bred with C57Bl6 mice for transmission of the recombined allele to the germline. Heterozygous F1 were bred with C57Bl6 mice to obtain mice harboring either a KI allele with normal nucleic acid sequence or an WT allele with normal nucleic acid sequence.

(b) Validation of recombination by Southern blot. Three DNA probes were used to ensure recombination of the allele in genomic DNA prior digested with BamHI. The *Nr3c1* 5' DNA probe detects the conKI allele, the *Neo* DNA probe detects the selection cassette and the *Flp* DNA probe detects the expression of the recombinase.

(c) Validation of recombination of the conKI allele by Southern blot. Three DNA probes were used to ensure recombination of the allele in genomic DNA prior digested with BamHI. The *Nr3c1* 5' DNA probe detects the conKI allele, the *Neo* DNA probe failed to detect the selection cassette removed in the previous generation, and the *CRE* DNA probe detects the expression of the recombinase.

(d) Typical result of genotyping by PCR of genomic DNA in 2% agarose gel electrophoresis. Specific bands are as follow: ConKI 491 bp, WT 265 bp, KI 318 bp.

(e) Schematic representation of the position of the genotyping primers in the various alleles.

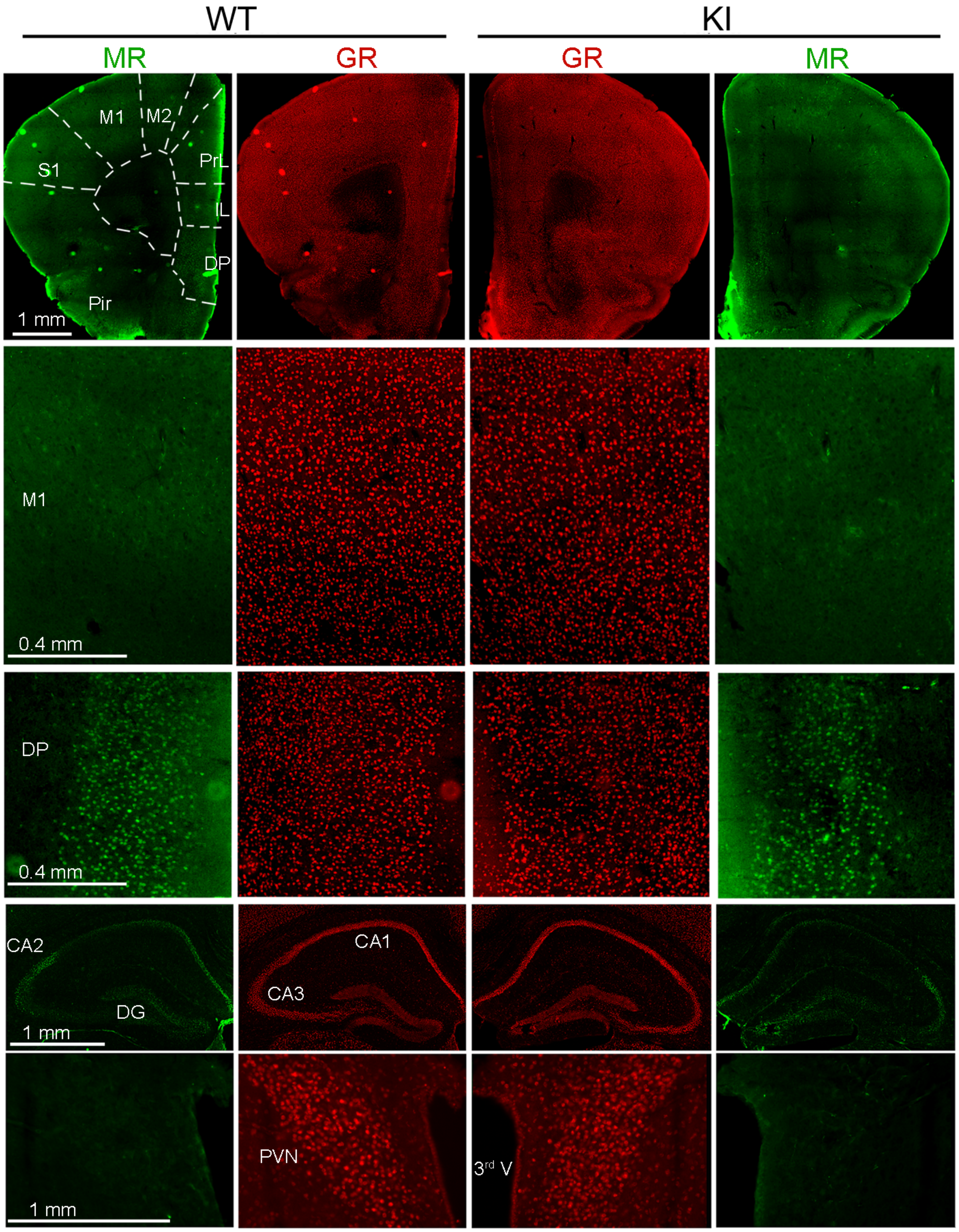


**Figure S4. Expression of glucocorticoid receptors (GR and MR) in brain of KI mice.** Detection of GR and MR in frontal cortex, hippocampus and hypothalamus reveals no difference of expression between WT and KI mice. M1-2 = motor area 1-2; S1 = sensory cortex area 1; Prl = prelimbic cortex; Il = infralimbic cortex; DP = dorsal peduncular cortex; Pir = piriform cortex; CA1-3 = cornus ammonis area 1-3; PVN = hypothalamic paraventricular nucleus.

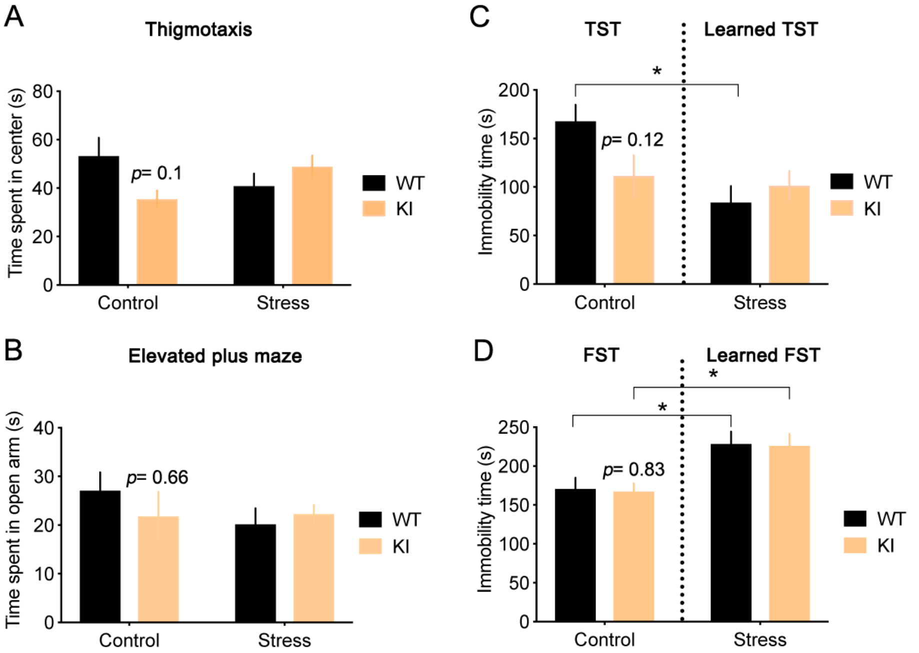


**Figure S5. No significant effects of genotype on the expression of anxiety and despair-like behaviors at 3 month of age**

(A) Time spent in the center arena of the open field. Means ±SEM of n=12 mice/ group, 2-way ANOVA: interaction of chronic unpredictable stress (for 3 weeks after weaning) and genotype $F_{1,44}$=5.7, $p$=0.0212 although genotype or stress alone had no effect.

(B) Time spent in the open arm of the elevated plus maze. No effect of chronic stress (for 3 weeks after weaning) and genotype alone or in interaction (means ±SEM of n=11 mice/ group).

(C) Time spent immobile in the tail suspension test (TST). Means ±SEM of n=11 mice/ group, 2-way ANOVA: Effect of habituation to the TST (Learned TST) in the stress group pre-exposed to this test once a week for 3 weeks $F_{1,40}$=6.9, $p$=0.0119.

(D) Time spent immobile in the forced swim test (FST). Means ±SEM of n=11 mice/ group, 2-way ANOVA: Effect of habituation to the FST (Learned FST) in the stress group pre-exposed to this test twice a week for 3 weeks $F_{1,40}$=16.94, $p$=0.0002.

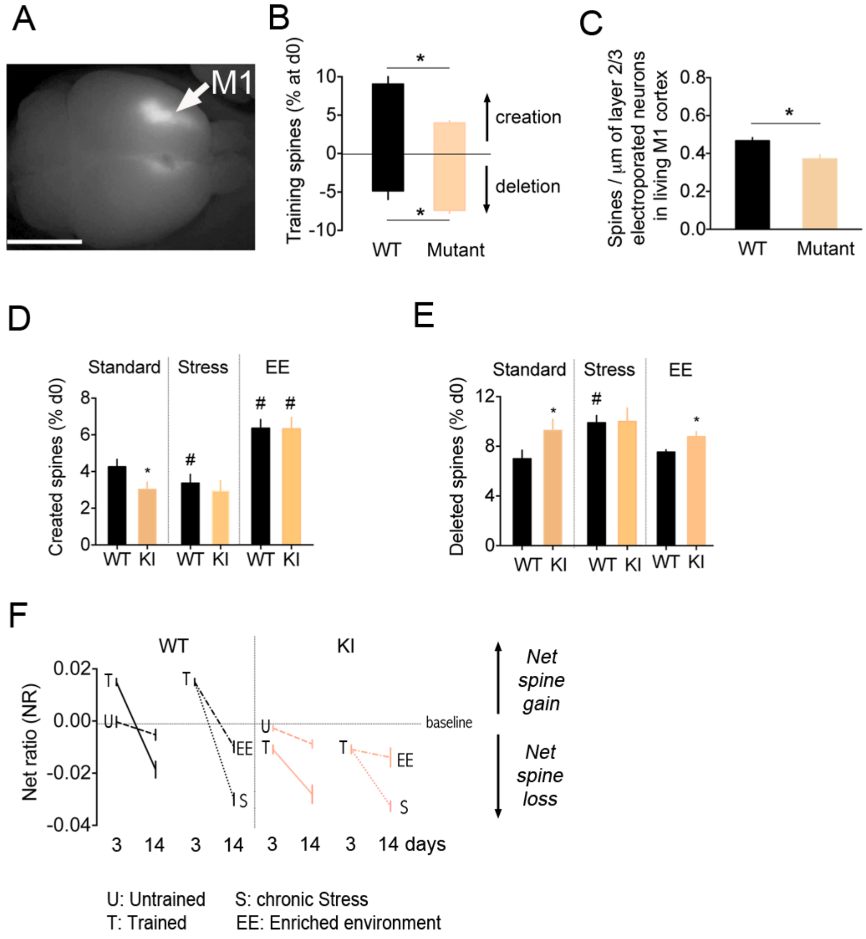


**Figure S6. Effect of GR-$PO_4$ deletion on experience-dependent spine dynamics.**

(A) *In utero* electroporation of GR constructs in excitatory NG neurons of M1 for substituting endogenous GR by recombinant GR-WT or mutant. Scale =4 mm.

(B) Remodeling of training spines in NG neurons electroporated with GR constructs. Means ±SEM of n=8 WT and 14 GR-mutant. Two-way ANOVA: Effect of GR-mutant $F_{(1,38)}$=25.1, $p$<0.0001, effect of spine remodeling $F_{(2,38)}$=225, $p$<0.0001, post-hoc Tukey test for spine creation *$p$=0.0008 and for spine deletion *$p$=0.031.

(C) Spine density at P35 in NG neurons of M1 electroporated with GR constructs. Means ±SEM of n=8 WT and 14 GR-mutant, unpaired t-test $t_{(11)}$=3.59 *$p$=0.0042.

(D) Spine formation (means ±SEM of n=8 WT standard, 8 KI standard, 7 WT stress, 7 KI stress and 7 WT EE, 7 KI EE mice). Two-way ANOVA: Effect of EE $F_{(1,26)}$=39.6, $p$<0.0001 post-hoc Tukey test in WT $^{\#}p$=0.027 and in KI $^{\#}p$<0.0001. Effect of chronic stress $F_{(1,26)}$=4.68, $p$=0.0039 post-hoc Tukey test $^{\#}p$=0.03.

(E) Spine deletion (means ±SEM of n=8 WT standard, 8 KI standard, 7 WT stress, 7 KI stress and 7 WT EE, 7 KI EE mice). Two-way ANOVA Effect of EE $F_{(1,26)}$=34.9, $p$<0.0001, post-hoc Tukey test in new spines *$p$=0.0002, in old spines *$p$=0.014. Effect of stress $F_{(1,26)}$=9.13, $p$=0.005 post-hoc Tukey test $^{\#}p$=0.0093.

(F) The fraction of spines gained subtracted of the spines lost day to day, were calculated as net ratio (NR) = ($N_{formed}$ - $N_{deleted}$)/(2 × $N_{total}$). Means ± SEM of n=6 WT and 6 KI untrained, 21 WT and 18 KI trained, 7 WT and 6 KI stress, 7 WT and 6 KI EE. Two-way ANOVA: effect of training in WT $F_{(1,36)}$=0.6 and in KI $F_{(1,32)}$=34.6, $p$<0.0001; effect of stress in WT $F_{(1,52)}$=9.7; $p$=0.0029 and in KI $F_{(1,44)}$=1; effect of EE in WT $F_{(1,52)}$=5.8; $p$=0.019 and in KI $F_{(1,44)}$=8.52; $p$=0.0055.

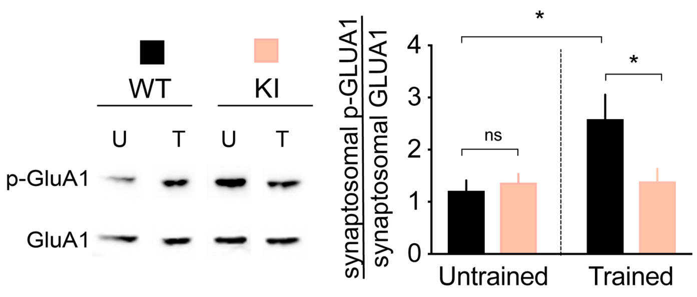


**Figure S7. Effect of GR-PO$_4$ deletion on training-induced GluA1 phosphorylation in motor cortex.**
(A) Synaptosomal GluA1-PO$_4$/GluA1 in M1 cortex. Means ±SEM, n=7 mice/group, 2-way ANOVA Effect of training $F_{1,24}$=5.7, $p$=0.025; training x genotype $F_{1,24}$=5.54, $p$=0.027 post-hoc Tukey test *$p$<0.05.

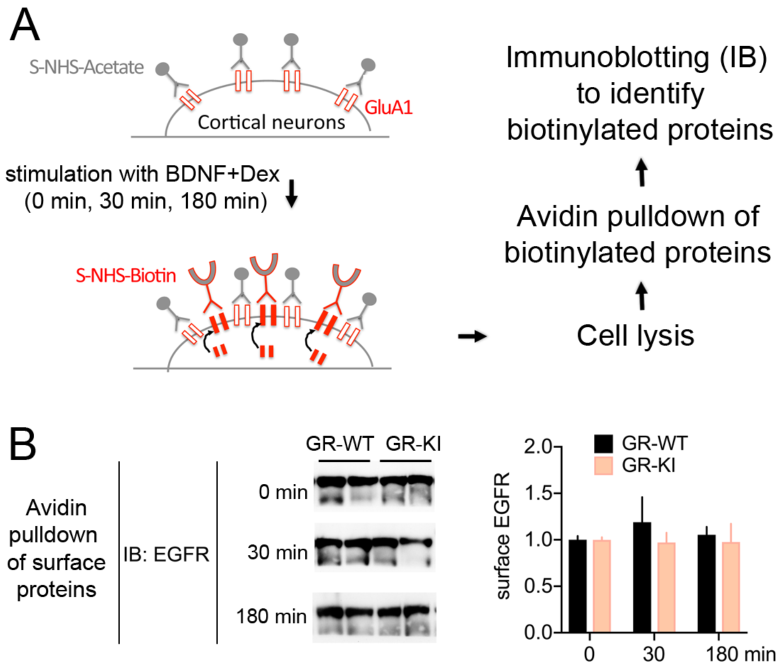


**Figure S8. No effect of GR-PO$_4$ deletion on the cell surface expression of EGFR.**
(A) Schematic representation of the procedure to isolate and monitor the dynamics of proteins insertion at the cell surface. Sulfo-NHS-acetate blocks all sites at the cell surface such that Sulfo-NHS-biotin can label newly inserted proteins. After cell lysis, avidin pulldown with magnetic beads allows for the purification of cell surface proteins that can be further identified by immunoblotting with specific antibodies.
(B) Avidin pulldown of biotinylated EGFR from neurons stimulated with 25 ng/ml BDNF and 1 μM Dex for the indicated time. Surface EGFR in cortical neurons expressing GR-WT or GR-KI constructs. Means ±SEM of n=5 WT and 5 KI/group, 2-way ANOVA: Effect of mutant $F_{1,24}$=0.67, $p$=0.4.

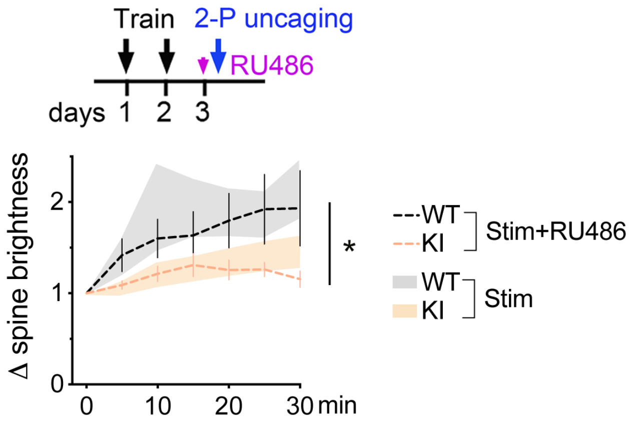


**Figure S9. No acute effect of RU486 on glutamate-evoked spine enlargement in motor cortex.**
Mice trained on the rotarod for 2 days were administered with RU486 (20 mg/kg, IP) 20 min before 2-P uncaging of glutamate at specific spine synapses in motor cortex. Means ±SEM of n=22 stim WT spines and 12 stim+RU486 WT spines, 40 stim KI spines and 18 stim+RU486 KI spines in 6 mice. Two-way ANOVA: Effect of RU486 $F_{3,593}=11.57$, $p<0.0001$; effect of time $F_{6,593}=4.64$, $p=0.0001$, post-hoc Tukey's test $*p<0.0127$.

### Antibodies

| Immunogen | Details | Source | Comments | Manufacturer |
|---|---|---|---|---|
| GR-$PO_4$ | S152 –P (S155-P in rat) | Rabbit polyclonal Affinity purified | Use at 1.5 mg/ml, epitope unmasking in 10 mM Tris PH=9, 1 mM EDTA, 0,05% tween for 20 min | Homemade (Lambert et al., 2013) |
| GR-$PO_4$ | S284-P (S287-P in rat) | Rabbit polyclonal Affinity purified | Use at 1.5 mg/ml, epitope unmasking in 10 mM Tris PH=9, 1 mM EDTA, 0,05% tween for 20 min | Homemade (Lambert et al., 2013) |
| GR-$PO_4$ | S246-P | Rabbit polyclonal Whole serum | Use at 1:500-2000 | Homemade (Ismaili and Garabedian, 2004) |
| GR-$PO_4$ | S232-P | Rabbit polyclonal Whole serum | Use at 1:1000 | Homemade (Ismaili and Garabedian, 2004) |
| GR-$PO_4$ | S224-P | Rabbit polyclonal Whole serum | Use at 1:1000 | Homemade (Ismaili and Garabedian, 2004) |
| GR | M20 | Rabbit polyclonal | Use at 1:400 | Santa Cruz Biotechnologies |
| Erk1/2 | | Rabbit polyclonal | Use at 1:1000 | Santa Cruz Biotechnologies |
| Parvalbumin | | Mouse polyclonal | Use at 1:3000 | Swant |
| Neurogranin | Ab5620 | Mouse polyclonal | Use at 1:5000, epitope unmasking in 10 mM Tris PH=9, 1 mM EDTA, 0,05% tween for 20 min | Millipore |
| GFP | | Chicken polyclonal | Use at 1:2000 | Abcam |
| RFP | | Rabbit polyclonal | Use at 1:1000 | Rockland |
| GR | BuGr2 | Mouse monoclonal | Use at 1:1000 | Calbiochem |
| PSD95 | clone K28/43 | Mouse monoclonal | Use at 1:1000 | NeuroMab |
| Synaptophysin | | Rabbit monoclonal | Use at 1:1000 | Life Technologies |
| EGFR | | Rabbit polyclonal | Use at 1:1000 | Cell Signaling Techologies |
| GLUA1 | | Rabbit polyclonal | Use at 1:1000 | Millipore |
| GLUA1-$PO_4$ | | Rabbit monoclonal | Use at 1:1000 | Millipore |
| c-FOS | | Mouse monoclonal | Use at 1:100, epitope unmasking in 10 mM Tris PH=9, 1 mM EDTA, 0,05% tween for 20 min | Santa Cruz Biotechnologies |
| c-FOS | #2250 | Rabbit monoclonal | Use at 1:1000 | Cell Signaling |
| AlexA Fluor conjugated secondary antibodies | | Rabbit monoclonal | Use at 1:2000 | ThermoFisher Scientific |

### Drugs

| Compound name | Effect | Working concentration | comments | Manufacturer |
|---|---|---|---|---|
| PMSF | Protease inhibitor | 100 ng/ml | In lysis buffer | Sigma |
| Aprotinin | Protease inhibitor | 100 nM | In lysis buffer | Sigma |
| Leupeptin | Protease inhibitor | 100 nM | In lysis buffer | Sigma |
| $Na_3VO_4$ | Tyr Phosphatase inhibitor | 1 mM | In lysis buffer | Sigma |
| NaF | Phosphatase inhibitor | 100 nM | In lysis buffer | Sigma |
| calyculin A | Phosphatase inhibitor | 10 nM | In lysis buffer | Sigma |
| BDNF | Neurotrophin | 25 ng/ml | TrkB agonist | Sigma |
| Dexamethasone | Synthetic glucocorticoid | 1 µM in vitro | GR agonist | Sigma |
| Bicuculline methiodide | GABAA receptor antagonist | 3.5 mM | Touch application | Sigma |
| MNI-glutamate | Caged-ligand Photoactivable | 20 mM | Topical application | Sigma |
| RU486 | GR antagonist | 20 mg/kg | intraperitoneal | Sigma |
| sulfo-NHS-acetate | Block cell surface proteins | 1.5 mg/ml | Use in TBS, 30 min at 4°C | Pierce |
| sulfo-NHS-LC-biotin | Labeling of surface proteins | 0.5 mg/ml | Use in TBS, 30 min at 4°C | Pierce |
| Glycine | Quench reactive sulfo-NHS moieties | 100 mM | Use in PBS, 20 min at 4°C | Sigma |
| Uridine | | 10 mM | Use with 5FU | Sigma |
| 5-fluoro-uridine | | 10 mM | Block cell proliferation | Sigma |
| B27 | Serum free | 2% (v/v) | Culture supplement | Life Technologies |

| Primers | |
|---|---|
| Gene | For genotyping of GR knockin mouse |
| Primer_01F | 5'-GCAGGCCGCTCAAGTGTTTTCT-3' |
| primer_02R | 5'-CACTGACCAACGAGAAACGATTAC-3' |
| primer_03F | 5'-GCAGGCAGAAGTGTGTTTAGC-3' |
| primer_04R | 5'-CAGTCATAGCCGAATAGCCTCTC-3' |
| primer_05F | 5'-CTGAGGGTGAAGACGCAGAAAC-3' |

**Table S1. List of antibodies, primers and chemical reagents used in study.**
